# Stimulus-Specific Modulation is Enabled by Differential Serotonin Receptor Expression

**DOI:** 10.1101/2023.06.21.546011

**Authors:** Julius Jonaitis, Mohd F.E.B. Mazri, Tyler R. Sizemore, Jacob D. Ralston, Farzaan Salman, Emma J. Fletcher, Danielle E. Matheny, Keshav L. Ramachandra, Andrew M. Dacks

**Affiliations:** Department of Biology, Life Sciences Building, West Virginia University, Morgantown, WV, 26506, USA; Department of Molecular, Cellular, and Developmental Biology, Yale Science Building, Yale University, New Haven, CT, 06520-8103, USA; Department of Neuroscience, West Virginia University, Morgantown, WV, 26506, USA

**Keywords:** Neuromodulation, Serotonin (5-HT), Olfaction, Connectomics, Drosophila

## Abstract

Neural networks must be able to flexibly process information under different conditions. To this end, networks frequently rely on uniform expression of modulatory receptors by distinct classes of neurons to fine tune the computations supported by each neuronal class. In this study, we explore the consequences of heterogeneous, rather than uniform, serotonin (5-HT) receptor expression within a cell class for olfactory processing in *Drosophila melanogaster*. Here, we demonstrate that two distinct populations of olfactory output neurons (projection neurons, PNs) display heterogeneous receptor co-expression of all 5-HT receptors. Moreover, the PN populations that express distinct 5-HT receptors innervate different combinations of glomeruli, implying that the effects of 5-HT on these PNs may vary with their odor tuning. Furthermore, connectomic analyses reveal that PN subsets with different receptor profiles have little convergence upon downstream synaptic partners. Finally, 5-HT differentially modulates the odor-evoked responses of PNs with distinct receptor expression profiles and odor tuning. Overall, this implies that heterogeneous modulatory receptor expression enables differential tuning of activity within a neuronal class depending on the odor scene to which individual neurons respond.

## Introduction

Within a sensory network, distinct neuronal classes influence the fidelity with which the nervous system can encode individual aspects of a given modality. For instance, inhibitory local interneurons (LNs) in the mammalian olfactory bulb and insect antennal lobe provide presynaptic inhibition of sensory afferents within the context of high stimulus intensity to avoid saturation of the olfactory system [1–7]. In this manner, one neuronal class impacts the encoding of a stimulus parameter within the context of ongoing sensory processing. Neuromodulation provides an elegant means by which to adjust the impact of individual network components, and the computations that they support, based on the ongoing behavioral state of the individual animal [8–10]. Neuromodulators usually activate several receptor proteins which allows a single signaling molecule to differentially affect cellular components that support distinct aspects of sensory coding. Often, a given neuronal class will express a consistent set of modulatory receptors, such that the entire population is affected in a similar manner and the neuronal computation that they exert can be uniformly altered [11]. For instance, a subset of GABAergic LNs in the antennal lobe (AL; first olfactory neuropil of *Drosophila*) express the 5-HT7 receptor allowing basal levels of 5-HT to affect the response gain modulation exerted by these LNs upon olfactory projection neurons (PNs) [12]. In an extreme example, 5-HT3 receptor expression is a distinguishing feature of a subset of GABAergic interneurons in the mammalian neocortex [13–16]. Although not g-protein coupled receptor, activation of the 5-HT3 receptor provides fast excitation to this heterogeneous population of interneurons [17,18] which are proposed to play a critical role in regulating the neuronal excitability based on behavioral state [14,18]. Thus, in these examples uniform expression of a 5-HT receptor allows fine tuning of the computation implemented by a population of neurons.

However, there are instances of heterogeneous expression of multiple receptor types for the same signaling molecule within a neuron class. Heterogeneous receptor expression suggests that a given neuromodulator can provide nuanced fine-tuning of response properties based on functional differences within that neuron class. For instance, all five insect 5-HT receptors are expressed within the populations of ventral and lateral-ventral projection neurons (v-PNs and lv-PNs respectively) in the AL of *Drosophila* [19]. Individual v-PNs and lv-PNs innervate different subregions of the olfactory system [20–23] raising the possibility that they may be modulated based on the individual volatile chemicals to which they respond, rather than uniformly across the population. Serotonin receptors differ in the binding affinity for 5-HT and the second-messenger systems to which they couple [24–26], implying that 5-HT may have a heterogeneous influence on the role of v-PNs and lv-PNs in olfactory processing. Here, we take advantage of the experimental tractability of *Drosophila* to explore the nature of within cell class modulatory receptor heterogeneity. We first demonstrate that there are many different patterns of 5-HT receptor co-expression amongst individual v-PNs and lv-PNs. Using stochastic labeling we demonstrate that v-PNs and lv-PNs expressing each 5-HT receptor innervate different but overlapping sets of glomeruli. We then identify driver lines for v-PNs with completely non-overlapping 5-HT receptor profiles and demonstrate that these populations have dissimilar odor tuning, sensitivity to 5-HT, and connectivity to third-order olfactory neurons within the LH. Overall, our results are consistent with a model in which 5-HT differentially modulates individual neurons within a cell class based on the stimuli to which they respond. Heterogeneous receptor co-expression therefore has the potential to expand the diversity of effects beyond those represented by the individual receptors.

## Results

The antennae and maxillary palps of *Drosophila* house thousands of olfactory sensory neurons (OSNs) that each express chemoreceptor proteins tuned to a set of volatile chemicals. OSNs that express the same chemoreceptive proteins converge upon a single glomerulus in the AL [27–29] where they synapse upon PNs. PNs integrate input from OSNs and local interneurons to generate responses that are relayed to the lateral horn (LH) and mushroom bodies (MBs), two second order olfactory neuropils associated with odor valence coding [30–32] and associative learning [33–37], respectively. PNs can be categorized based on their glomerular innervation, the tract by which their axons leave the AL, the cell body cluster to which they belong and their transmitter content. There are ∼127 mostly uniglomerular cholinergic PNs deriving from anterodorsal and lateral cell body clusters surrounding the AL and these project along the medial AL tract (mALT) to innervate first the MB and then the LH [22,38]. A third, ventrally located cell body cluster contains predominantly multiglomerular PNs. These PNs are subdivided into cholinergic lv-PNs that project along the mALT to the superior intermediate protocerebrum then the LH, and GABAergic v-PNs that project along the medial-lateral AL tract (mlALT) directly to the LH [22,39]. Based on a whole brain EM dataset [40], initial estimates place the counts of v-PNs and lv-PNs at just over 70 PNs of each type per AL [22], although this might be an underestimation [41]. Altogether, the individual innervation patterns of PNs within the LH is consistent across flies, establishing a zonal map of broad categories such as food or pheromone related odorants [21,42–49]. The GABAergic v-PNs are more broadly tuned than the uniglomerular cholinergic PNs [43], integrating interglomerular synaptic input from ORNs and excitatory PNs that provide output locally within the AL [43]. The v-PNs provide feedforward inhibition to the LH [21,50,51] that refines the spatiotemporal patterns of odor-evoked activity of different LH neuron classes [21,31,43,51–53] and has been implicated in playing a role in odor discrimination [43,50], valence coding [21,51], and habituation [54]. Finally, the AL, LH, mushroom body calyces and several third order olfactory regions are innervated by a pair of serotonergic neurons called the CSDns (contralaterally-projecting serotonergic deutocerebral neurons) [55–59] which have extensive synaptic connectivity with each of the principal AL cell classes [60,61].

### v-PNs and lv-PNs display heterogeneous 5-HT receptor co-expression

While v-PNs and lv-PNs collectively express all five 5-HT receptors [19], it remains unclear if they co-express 5-HT receptors and if there are any common 5-HT receptor co-expression profiles among v-PNs. To resolve this, we took an intersectional approach leveraging a set of 5-HT receptor MiMIC T2A-Gal4 lines [62] and LexA knock-in lines [63] to make pairwise comparisons of receptor expression. Only the 5-HT2B and 5-HT7 LexA knockin lines had 100% overlap in their v-PN and lv-PN expression patterns with their respective MiMIC T2A-Gal4 driver line (**Figure 1A, Figure S1)**, so we restricted our comparisons to the 5-HT2B and 5-HT7 knock-in LexA lines with the five MiMIC 5-HT receptor T2A lines (**Figure 1A-C)**. For both the v-PNs and lv-PNs, most of the five 5-HT receptors were co-expressed, except for the 5-HT2A receptor which was not co-expressed with either the 5-HT2B or 5-HT7 receptors (**Figure 1D, Figure S1D-U)**. The v-PNs tended to have slightly higher degree of co-expression of the 5-HT1A receptor with both the 5-HT2B and 5-HT7 receptors (∼30% each) relative to the lv-PNs, while the pattern was reversed for co-expression of the 5-HT1B receptor in lv-PNs (∼25% for the 5-HT2B and ∼40% for the 5-HT7). Thus, 5-HT likely exerts a heterogeneous impact on the total population of v-PNs and lv-PNs, greater than could be achieved with either non-overlapping patterns of 5-HT receptor expression or a uniform pattern of expression.

**Fig. 1.**
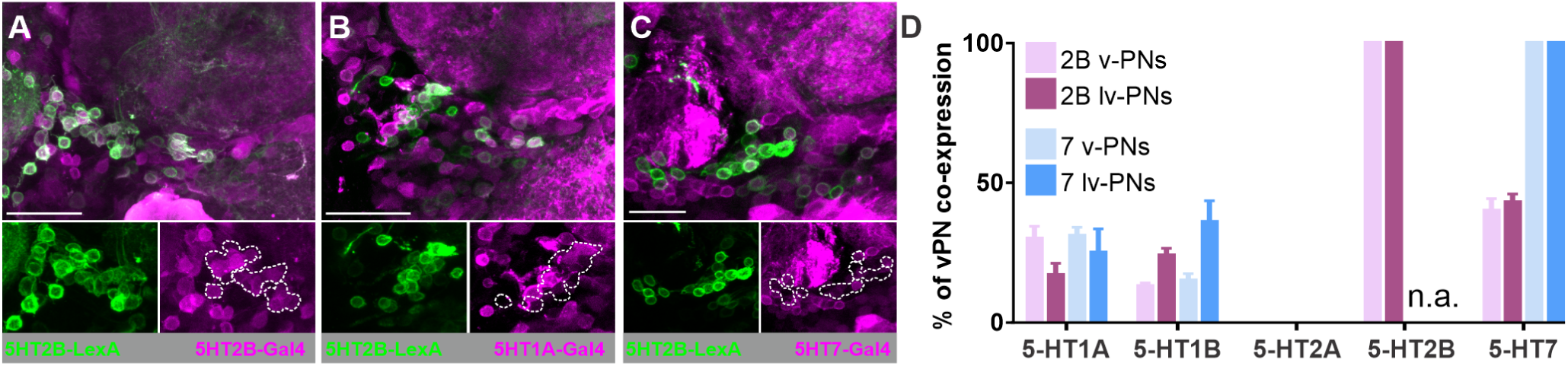
As a population v-PNs express a diverse combination of serotonin receptors. **A)** Example v-PN cell clusters demonstrating degree of overlap of expression the 5-HT2B LexA with **A)** the 5-HT2B-T2A-Gal4, **B)** the 5-HT1A-T2A-Gal4 and **C)** the 5-HT7-T2A-Gal4. **D)** Percent of overlap between v-PNs expressed in either the 5-HT2B-LexA (blue) or 5-HT7-LexA (pink) with v-PNs in the T2A-Gal4 lines for the other 5-HT receptors.

The diverse set of co-expression patterns displayed by v-PNs and lv-PNs could still have a uniform effect on the role of these populations in olfactory coding if v-PNs and lv-PNs expressing each 5-HT receptor innervate the same AL glomeruli and sub-regions within the LH. To determine if differential 5-HT receptor expression by v-PNs and lv-PNs also correlated with differences in odor tuning, we used the multi-color flip-out (MCFO) technique [64] to stochastically label v-PNs and lv-PNs within the 5-HT receptor MiMIC T2A-Gal4 lines and scored the glomeruli that they innervated (**Figure 2A-C)**. We note that this approach likely highlights v-PNs and lv-PNs with the highest levels of expression of a given 5-HT receptor. Although this likely represents a biased sampling, it is still informative about the odor information sampled by v-PNs and lv-PNs that express a given 5-HT receptor. Across over 200 ALs (**Figure 2D)** we were more likely to observe lv-PNs for the 5-HT1A (70% of samples), 5-HT1B (94% of samples), 5-HT2A (100% of samples) and 5-HT2B (96% of samples) T2A-Gal4 lines, while we were more likely to observe v-PNs for the 5-HT7 T2A-Gal4 line (96% of samples). Displaying the percent of samples in which a given glomerulus was innervated by a v-PN or lv-PN flipped out in a given 5-HT receptor T2A line illustrates that the 5-HT1A, 5-HT1B, 5-HT2A and 5-HT2B were more reliably observed innervating (or not innervating) a given set of glomeruli, whereas the v-PNs expressing the 5-HT7 receptor were more variably observed, but innervated almost every glomerulus (**Figure 2E)**. This implies that the patterns of glomerular innervation for the 5-HT7 receptor were the most dissimilar from the other four receptors. Using hierarchical clustering to group glomerular innervation patterns from individual brains revealed that v-PNs and lv-PNs from each 5-HT receptor MiMIC T2A-Gal4 generally do not segregate into their own unique clusters **(Figure S2)**. Instead, two clusters formed predominantly composed of samples from the 5-HT7 T2A-Gal4 line and three clusters that included samples derived from the other four 5-HT receptor T2A-Gal4 lines. Overall, this approach suggests that v-PNs and lv-PNs with distinct 5-HT receptor expression patterns innervate different combinations of glomeruli. Given this framework, 5-HT is poised to modulate the v-PN and lv-PN populations in a stimulus specific manner.

**Fig. 2.**
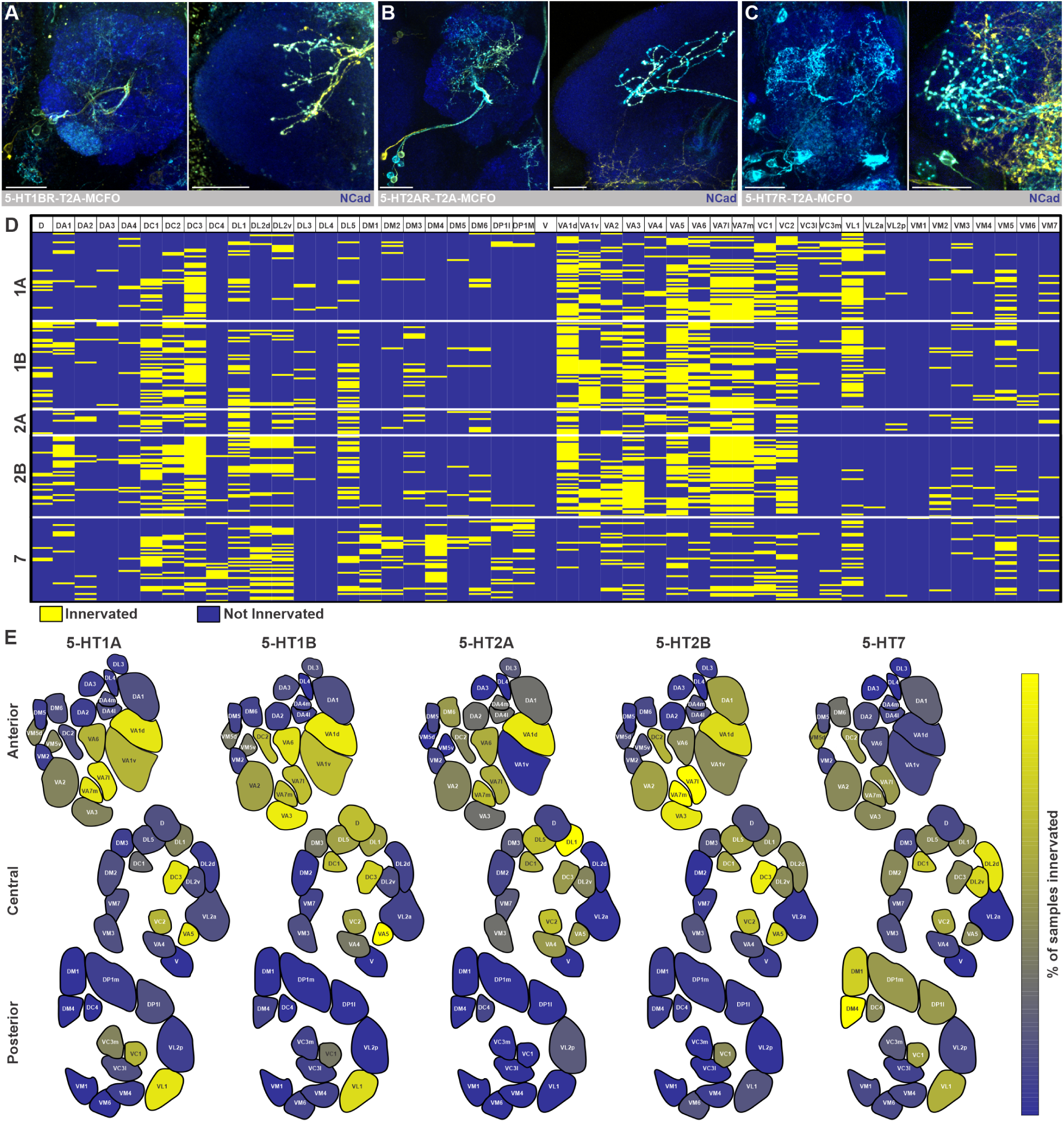
Population analysis of v-PNs expressing each 5-HT receptor. **A)** Example image of v-PN morphology within the AL (left panel) and LH (right panel) revealed by MCFO technique using the **A)** 5-HT1A-T2A-Gal4 (lv-PN), **B)** 5-HT2A-T2A-Gal4 (lv-PN) and **C)** 5-HT7-T2A-Gal4 (v-PN). **D)** Glomerular innervation patterns of v-PNs stochastically labeled via MCFO using each of the 5-HT receptor T2A-Gal4 lines. Each row represents the glomeruli innervated by v-PNs in a single AL with yellow indicating innervation of a given glomerulus and dark blue indicating no innervation. **E)** Glomerular maps depicting percent of samples in which a given glomerulus was innervated by v-PNs stochastically labeled using each of the 5-HT receptor T2A-Gal4 lines.

### Differential stimulus-specific modulation of v-PNs is enabled by distinct 5-HT receptor expression

To test the hypothesis that 5-HT differentially modulates v-PNs with distinct odor tuning, we sought to identify driver lines in which only a small number of v-PNs are included, all of which express the same 5-HT receptor profile. After an initial screen of the FlyLight database [65] to identify driver lines with expression of only a few v-PNs within the AL, we identified R24H08 (**Figure 3A)** and R86G06 (**Figure 3B)** as driver lines in which all v-PNs have the same 5-HT receptor expression profile. Unfortunately, we were unable to find lv-PN driver lines that met these criteria. R24H08 had some additional neurons expressed within the LH, so we created a split-Gal4 line **(see Methods)** to restrict expression solely to the ∼5 v-PNs in this line **(Figure S3A, B)**. R24H08 v-PNs densely innervate VC2, DC3 and VL2a (**Figure 3C)**, whereas the ∼5 v-PNs in R86G06 sparsely innervate over 20 glomeruli (**Figure 3D)**, with only VC2 overlapped with the glomeruli innervated by R24H08. The v-PNs expressed in R24H08 were essentially uniglomerular (except for some sparse branching in other glomeruli) resembling the “mlPN1” type neurons described by [39], and thus differing morphologically from v-PNs in R86G06 which were multiglomerular similar to the “mlPN2” type neurons [39]. These two sets of v-PNs also innervate distinct regions of the LH, with the R24H08 v-PNs innervating the ventromedial LH **(Figure 3A)** and the R86G06 v-PNs innervating the dorsomedial LH (**Figure 3B)**. Thus, these two sets of v-PNs innervated mostly non-overlapping sets of AL glomeruli (with the exception of VC2) and LH regions. Both sets of v-PNs expressed GAD1 indicating that they provide feedforward inhibition to the LH **(Figure S3C-D)**. GABAergic v-PNs have almost exclusively been studied using the Mz699 driver line [21,39,43,50–52], however the intersection of Mz699-Gal4 with each v-PN LexA revealed no overlap between R24H08-LexA or R86G06-LexA **(Figure S3E-H)**. Finally, we used the 5-HT receptor T2A-Gal4 lines [62] in combination with the R24H08 and R86G06 LexA lines to screen each v-PN subset for 5-HT receptor expression. This approach revealed that the R24H08 v-PNs exclusively expressed the 5-HT1A and 5-HT1B receptors (**Figure 3E-F, 3H & Figure S3I-M)**, whereas the R86G06 v-PNs only expressed the 5-HT7 receptor (**Figure 3G-H & Figure S3N-R)**.

**Fig. 3.**
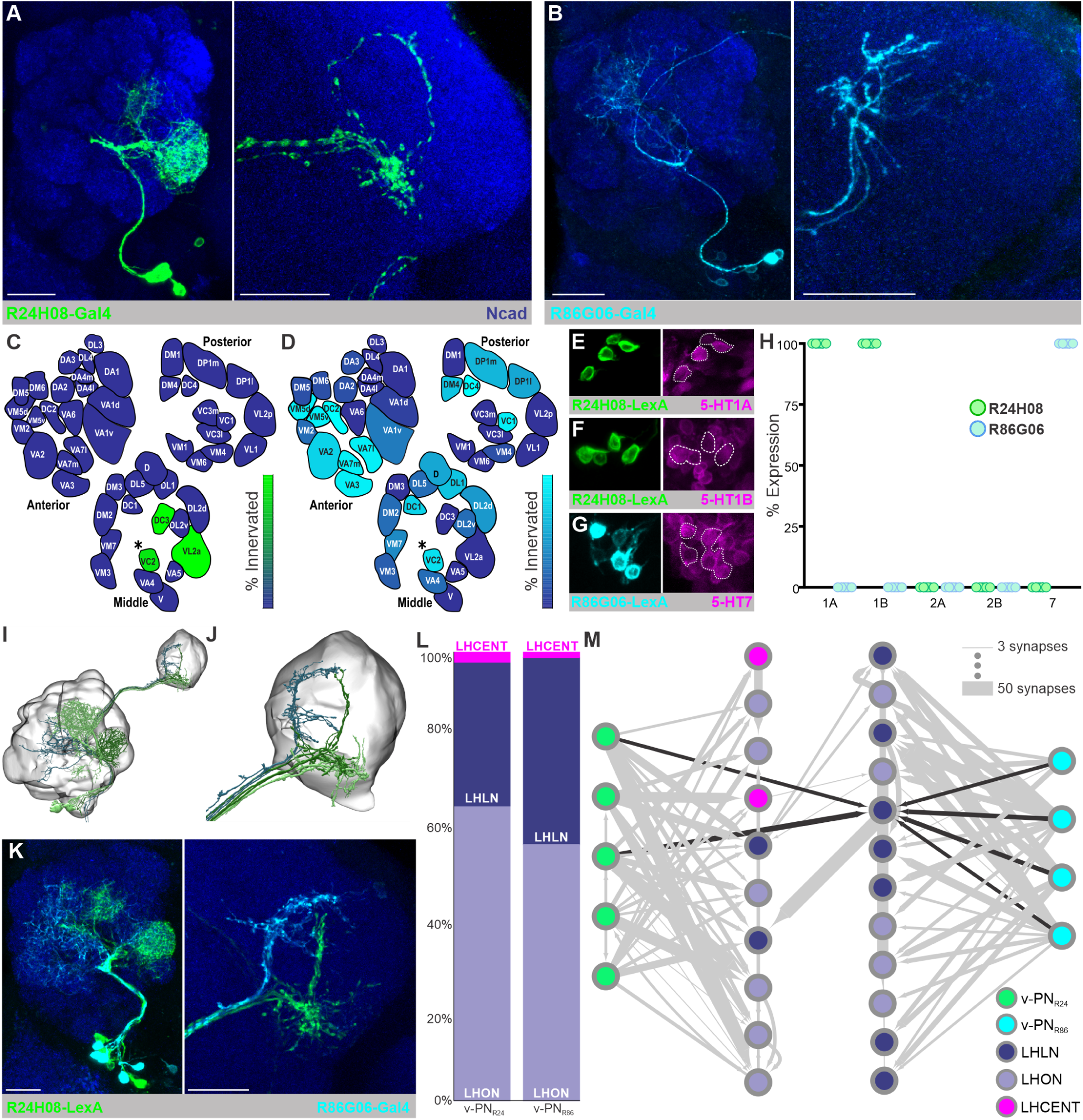
v-PNs innervating separate sub-regions of the olfactory system express different 5-HT receptor combinations and have differential odor-tuning. **A)** v-PNs expressing different serotonin receptors project to distinct regions of the olfactory system. **A)** Morphology of the v-PNs within the R24H08-Gal4 (green) within the AL and LH. Immunolabeling for N-Cadherin (blue) highlights neuropil. **B)** Morphology of the v-PNs within the R86G06-Gal4 (cyan) within the AL and LH. **C)** Map of glomeruli innervated by R24H08 v-PNs. **D)** Map of glomeruli innervated by R86G06 v-PNs. **E)** R24H08 v-PNs overlap with the 5-HT1A-T2A-Gal4. **F)** R24H08 v-PNs overlap with the 5-HT1B-T2A-Gal4. **G)** R86G06 v-PNs overlap with the 5-HT7-T2A-Gal4. **H)** Percent of v-PNs within the R24H08 (green) and R86G06 (blue) LexAs expressed within each 5-HT receptor T2A-Gal4 line. **I)** Reconstruction of the v-PNs from the Hemibrain dataset that most closely resemble the R24H08 (v-PN_R24_; green) and R86G06 (v-PN_R86_; dark blue) driver lines within AL and LH volumes. R24H08 v-PNs are shown in two shades of green to highlight the v-PNs predominantly innervating the VL2a glomerulus (dark green) and the DC3 glomerulus (light green). **J)** Reconstruction of the axon terminals of the two v-PN groups within the LH. **K)** overlap of AL (left) and LH (right) projections of the R24H08 (green) and R86G06 (cyan) v-PNs. Immunolabeling for N-Cadherin (blue) highlights neuropil. **L)** Synapse fractions of the downstream LH neurons targeted by the candidate v-PNs from the R24H08 and R86G06 driver lines. LH output neurons (LHONs; lavendar), LH local neurons (LHLNs; purple), LH centrifugal neurons (LHCENTs; pink). **M)** Graph plot of the downstream synaptic targets of the v-PN_R24_ (green) and v-PN_R86_ (cyan) shows very little convergence, except for a single LHLN that is targeted by two v-PN_R24_ and all four v-PN_R86_ (black arrows). All scale bars = 20um.

Next, we sought to determine the degree to which these two v-PN subsets converge upon the same downstream targets within the lateral horn. Theoretically, third-order olfactory neurons within the lateral horn could integrate synaptic input from v-PNs differentially modulated by 5-HT, allowing serotonergic modulation to shift which odor-evoked responses have the greatest impact on a common partner. Alternatively, v-PNs that express different 5-HT receptors could synapse on non-overlapping populations, so that the impact of serotonergic modulation on each set of v-PNs remains separate. To address this question, we turned to the hemibrain dataset [66], a nanoscale resolution electron microscopy (EM) volume that includes the right AL and LH of a female *Drosophila*. By querying v-PNs in hemibrain based on the AL glomerular innervation patterns observed for the R24H08 and R86G06 driver lines, we identified five and four v-PNs within the EM dataset that closely resembled the v-PNs expressed by the R24H08-Gal4 and the R86G06-Gal4 lines, respectively (referred to as v-PN_R24_ and v-PN_R86_) (**Figure 3I & J)**. In particular, the branching patterns of the v-PNs within the LH of the EM dataset (**Figure 3J)** were very similar to that observed from light microscopy data (**Figure 3K)**. To determine if these two groups of v-PNs synapse upon similar neuronal demographics within the LH, we queried all of the downstream partners of the nine v-PNs and classified each downstream partner within the LH as an LH output neuron (LHON), LH centrifugal neuron (LHCENT), or LH local neuron (LHLN) based on the cell body tract to which they belong, and the relative distribution of presynaptic and post-synaptic sites within the LH and other brain regions, as described previously [67,68]. The broad demographics of the LH neurons targeted by each v-PN group was very similar (**Figure 3L)**, with both primarily targeting LHONs, a smaller proportion of LHLNs and very few LHCENTs. We then searched for all downstream partners that receive convergent synaptic input from two or more of the nine total v-PNs (regardless of the driver line that they resemble). Of the 22 total lateral horn neurons that receive convergent input from two or more of the queried v-PNs, only 1 LHLN was downstream of both v-PN_R24_ and v-PN_R86_ (**Figure 3M)**. Thus, while the two groups of v-PNs target similar neuron-class demographics, there was almost no convergence in the downstream neuronal populations targeted by v-PNs that express different 5-HT receptors (**Figure 3M)**. This implies that at the level of the LH the impact of 5-HT on v-PNs that express different 5-HT receptors remains segregated.

We next compared the odor tuning of the v-PNs expressed by the R24H08 and R86G06 driver lines. We selected a panel of odors based upon previous reports of odor-evoked responses of OSNs innervating the glomeruli also innervated by each v-PN driver line [69–73] and odor blends such as apple cider vinegar (ACV) [74] and orange peels [75]. We then used calcium imaging of LH branches to visualize odorevoked responses of each line. Using this approach, we identified benzaldehyde as an odorant that activated both sets of v-PNs, farnesol as an odorant that only activated R24H08 v-PNs and 1-octen-3-ol and 1-hexanol as odorants that preferentially activated R86G06 (**Figure 4A & B)**. Thus, these v-PNs differ in the regions of the olfactory system that they innervate, the 5-HT receptors they express and their odor tuning.

**Fig. 4.**
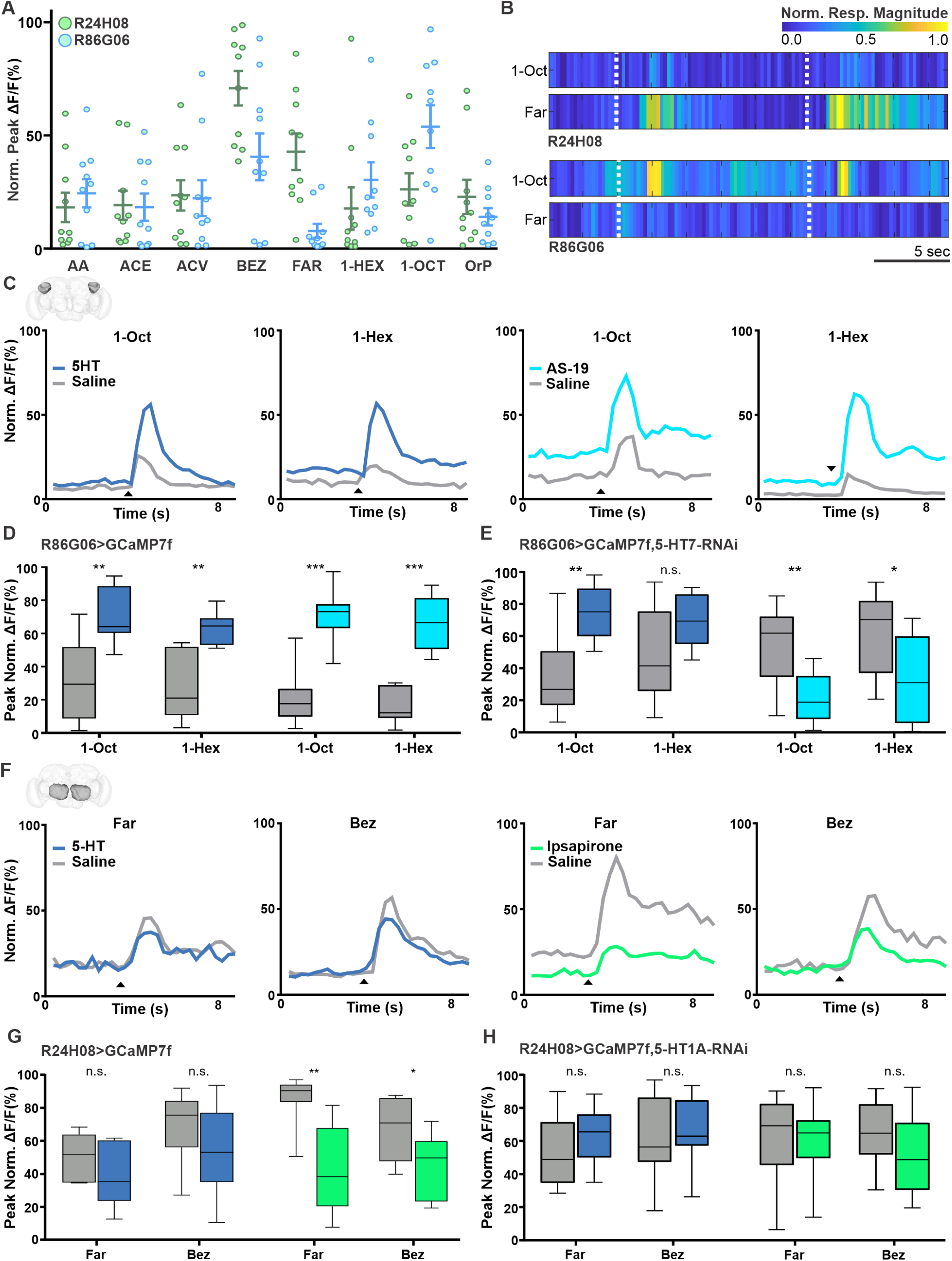
Serotonin differentially modulates odor-evoked responses of v-PNs. **A)** Calcium imaging responses (peak %ΔF/F) of R24H08 (green) and R86G06 (cyan) v-PNs to a panel of odors. AA; acetoin acetate, ACE; acetophenone, ACV; apple cider vinegar, BEZ; benzaldehyde, FAR; farnesol, 1-HEX; 1-hexanol, 1-OCT; 1-octen-3-ol, OrP; orange peel (n=10). **B)** R24H08 v-PNs respond to farnesol, but not 1-octen-3-ol, while R86G06 v-PNs respond to 1-octen-3-ol, but not farnesol. **C)** Average responses of R86G06 v-PNs to 1-octen-3-ol or 1-hexanol before (gray traces) and after application of 5-HT (dark blue) or a 5-HT7 receptor agonist (AS-19; cyan). Arrowhead indicates time of odor application. **D)** 5-HT and AS-19 enhance odor-evoked responses of R86G06 v-PNs. 5-HT, n=10 (1-Oct), p=0.0057; n=7 (1-Hex), p=0.0053; AS-19, n=10 (1-Oct), p=0.0003; n=8 (1-Hex), p=0.0002; 3 repeated stimulations, significance tested using paired samples T-test. **E)** RNAi knockdown of the 5-HT7 receptor in R86G06 v-PNs results in a loss of 5-HT induced enhancement of 1-hexanol responses and causes AS-19 to suppress R86G06 v-PN responses. 5-HT, n=10 (1-Oct), p=0.0051; n=9 (1-Hex) n.s.; AS-19, n=11 (1-Oct), p=0.0041; n=12 (1-Hex) p=0.0323; 3 repeated stimulations, significance tested using paired sample T-test. All recordings were made from R86G06 v-PN axon terminals in the lateral horn. **F)** Average responses of R24H08 v-PNs to farnesol (Far) or benzaldehyde (Bez) before (gray traces) and after application of 5-HT (blue) or a 5-HT1A/1B receptor agonist (Ipsapirone; green). Arrowhead indicates time of odor application. **G)** 5-HT does not significantly affect odor-evoked responses of R24H08 v-PNs, while ipsapirone reduces responses. 5-HT, n= 7 (Far), n.s.; n=11 (Bez), n.s.; Ipsapirone, n=7 (Far), p=0.0016; n=11 (Bez), 3 repeated stimulations, significance tested using paired sample T-test. **H)** RNAi knockdown of the 5-HT1A receptor in R24H08 v-PNs results in a loss of the ipsapirone induced reduction of odor-evoked responses. 5-HT, n=8 (Far), n.s.; n=9 (Bez), n.s.; Ipsapirone, n=14 (Far), n.s.; n=13 (Bez), n.s.; 3 repeated stimulations, significance tested using paired sample T-test. All recordings were made from R24H08 v-PN processes in the AL.

Having identified odorants that strongly activated v-PNs within each driver line, we then tested whether 5-HT differentially impacts their odor-evoked responses. We began by visualizing GCaMP transients in the axon terminals of each set of v-PNs within the LH in response to either farnesol and benzaldehyde, or 1-octen-3-ol and 1-hexanol, for the v-PNs in R24H08 and R86G06 respectively. We then bath applied 5-HT, metitepine (an antagonist of all five *Drosophila* 5-HT receptors; [24]), 5-HT receptor specific agonists (ipsapirone for the 5-HT1A and 5-HT1B receptors, and AS-19 for the 5-HT7 receptor) and 5-HT receptor specific antagonists (WAY100635 for the 5-HT1A and 5-HT1B receptors, and SB269970 for the 5-HT7 receptor). The odor-evoked responses of R86G06 v-PNs to 1-octen-3-ol and 1-hexanol were significantly enhanced by both 5-HT and the 5-HT7 receptor agonist AS-19 (**Figure 4C & D)**. However, neither metitepine, nor the 5-HT7 receptor antagonist SB269970 affected the odor-evoked responses of R86G06 v-PNs suggesting that under the conditions in which these recordings were performed, basal 5-HT levels do not impact their activity **(Figure S4A & B)**. Driving a 5-HT7 receptor RNAi in R86G06 v-PNs eliminated the 5-HT induced enhancement of responses to 1-hexanol, but not 1-octen-3-ol (**Figure 4E)**. This may indicate that the enhancement of R86G06 v-PN responses to 1-octen-3-ol is polysynaptic in nature, whereas the enhancement of R86G06 v-PN responses to 1-hexanol is due to direct modulation. The odorant-dependent variability in the impact of 5-HT7 RNAi in R86G06 v-PNs was consistent with similar observations for cholinergic PNs in *Drosophila* [76] and other insects [77]. Furthermore, the serotonergic neurons innervating the olfactory system of *Drosophila* indirectly impact multiglomerular PNs in larvae by modulating local interneurons in larvae [78]. Surprisingly, R86G06 v-PN responses to 1-octen-3-ol and 1-hexanol were both significantly reduced by AS-19 when R86G06 expressed the 5-HT7 RNAi. This suggests that AS-19 both directly modulates R86G06 v-PNs and enhances inhibitory input to the R86G06 v-PNs which is revealed once the direct modulation is blocked.

Since the 5-HT1A and 5-HT1B receptors are negatively coupled to adenylyl cyclase, we expected that the odor-evoked responses of R24H08 v-PNs would be reduced by 5-HT. However, 5-HT and ipsapirone (a 5-HT1A and 5-HT1B agonist) only caused a slight, non-significant reduction in odor-evoked responses of R24H08 v-PN processes recorded in the LH **(Figure S4C & D)**. Furthermore, neither the general 5-HT receptor antagonist metitepine nor the 5-HT1A receptor antagonist WAY100635 affected the magnitude of odor-evoked responses recorded from R24H08 v-PN processes within the LH **(Figure S4E & F)**. One possible explanation for this apparent lack of effect was that bath application of pharmacological agents induced a transient effect on R24H08 v-PNs that dissipated between odor trials. However, bath application of ipsapirone for 8 minutes did not cause a noticeable change to the baseline fluorescence of R24H08 v-PNs, even though AS-19 caused an increased baseline fluorescence for R86G06 v-PNs **(Figure S4G)**. Another possible explanation for the lack of impact of 5-HT receptor pharmacology on the odor-evoked responses of R24H08 v-PNs could be that the impact of 5-HT may be localized to the v-PN processes within the AL, rather than the LH. We therefore repeated the experiments above while imaging in the AL and found that ipsapirone significantly reduced the responses of R24H08 v-PNs to farnesol and benzaldehyde recorded within the AL (**Figure 4F & G)**. Furthermore, expression of 5-HT1A RNAi in the R24H08 v-PNs eliminated this reduction (**Figure 4H)** implying that it was due to direct receptor agonism by ipsapirone.

## Discussion

Each neuronal class within any sensory network supports distinct computations, however the relative importance of each computation can be modified based on recent network activity or the current physiological needs of the individual animal. In the olfactory system of *Drosophila*, serotonergic neurons interact with most principal neuron classes [59,60] to impact olfactory processing and behavior [61,79–81]. To this end, the diverse suite of serotonin receptors expressed by each neuronal class provides precise regulation of distinct computations so that network activity can be contextually optimized [82–84]. Frequently neurons within a given neuronal class will uniformly express one or a few modulatory receptors enabling a neuromodulator to adjust the specific computation. For instance, within the lamina of the *Drosophila* optic lobe, L2 laminar monopolar cells express the 5-HT2B receptor, whereas medullar T1 cells express the 5-HT1A and 5-HT1B receptors [85]. Since T1 and L2 cells play distinct roles in visual processing [86], this suggests that 5-HT differentially affects the computations supported by each cell class. In this study, we demonstrate that even within a neuronal class there can exist combinatorial receptor expression allowing a single modulator to exert its effects in a stimulus-specific manner.

A diverse set of receptors provides the nervous system with a great deal of flexibility for regulating the activity of individual cellular components within a network. A single signaling molecule such as 5-HT can have a range of effects as 5-HT receptors differ in their threshold for activation, time course of action, and the valence of their impact on neuronal activity. The half-effective concentrations of 5-HT for *Drosophila* 5-HT receptors differs over a range of two-orders of magnitude [24]. Collectively 5-HT receptors couple with G_s_, G_q_, and G_i_ (reviewed in [25] and [26]) allowing for activation of a range of transduction mechanisms with different degrees of potential amplification. We report that not only are all five 5-HT receptors expressed by v-PNs, but that there is extensive co-expression of 5-HT receptors (**Figure 1)**. Although we could not test all pairwise combinations, v-PNs and lv-PNs have some degree of co-expression for receptors that couple to different second messenger pathways. For instance, the 5-HT2B receptor (which couples to IP_3_ pathways) is co-expressed with both 5-HT1 receptor subtypes and the 5-HT7 receptor (all three of which impact adenylyl cyclase). Furthermore, co-expression of the 5-HT7 receptor (which positively couples to adenylyl cyclase via G_s_) and the 5-HT1 receptors (which negatively couple to adenylyl cyclase via G_i_) indicates that there is integration within even a single second-messenger pathway. Co-expression effectively increases the potential effects of 5-HT from the singular actions of each individual receptor, to the wider range of effects endowed by combinatorial patterns of receptor expression. There is also the possibility that 5-HT may have synergistic effects on v-PNs or lv-PNs due to the formation of 5-HT receptor heterodimers (reviewed in [87] and [88]). For instance, 5-HT1A receptor induced GPCR inward rectifying potassium (GIRK) channel currents are reduced when cells co-express the 5-HT7 receptor, even though the 5-HT7 receptor does not itself target GIRK [89]. Thus, co-expression of 5-HT receptors can result in interactions between receptors and convergence of downstream transduction pathways.

Our approach only allowed us to determine the neurons that express a given 5-HT receptor, but not where that receptor is trafficked. Compartment specific receptor trafficking may explain why the effects of 5-HT1A/B agonism were more potent for R24H08 v-PN processes within the AL relative to the LH (**Figure 4 & Figure S4)**. With the advances in conditional protein tagging methods [90], it will be exciting to determine if neurons that span multiple networks also differentially traffic modulatory receptors. In the case of 5-HT within the olfactory system of *Drosophila*, this is especially important as the branches of the CSDns within the AL are inhibited by odors in a concentration dependent, odor-independent manner, whereas within the LH they are excited in a concentration independent, odor-dependent manner [91]. Furthermore, the CSDns have more synapses upon the processes of v-PNs and lv-PNs in the AL than in the LH [60], although the CSDns may implement paracrine signaling in addition to synaptic transmission [79]. Differential receptor trafficking could therefore impact the context in which 5-HT affects a single postsynaptic neuron within the olfactory system. Compartment specific modulation appears to be common within the mushroom bodies of *Drosophila* [92–94] and could conceivably occur for v-PNs in the LH as well.

Both v-PNs and lv-PNs appear to play a role in supporting innate odor-driven behavior. In a recent study on directional processing of odor information, lv-PNs responsive to the pheromone cis-vaccenyl acetate were shown to play a critical role in promoting copulation rate and male-male aggression [95]. The GABAergic v-PNs have received far more attention for their role in refining odor representations in the *Drosophila* LH and appear to play an important role in odor decorrelation in support of innate odor-guided behavior [21,50,51,96]. Although not an example of feedforward inhibition, GABAergic cortical projections to olfactory bulb decorrelate tufted cell odor-evoked responses in mammals [97] similar to the impact of v-PNs on glutamatergic LH neurons in *Drosophila* [96]. Furthermore, inhibition of mammalian GABAergic corticobulbar projections [97] and inhibition of the neurons expressed in the Mz699-Gal4 line in *Drosophila* [50] reduces odor discrimination in behavioral assays, demonstrating a role for long-range inhibitory projections in fine odor coding. Glutamatergic corticobulbar feedback on the other hand, decorrelates mitral cell odor representations via disynaptic inhibition [98,99] and thus may play a complementary role to GABAergic corticobulbar feedback in mammals. It remains to be seen however, if the cholinergic lv-PNs play a similarly complementary role with the GABAergic v-PNs within the LH. Regardless of their relative roles, our data suggests that small subsets of v-PNs and lv-PNs that have distinct odor-tuning express different combinations of 5-HT receptors allowing their influence on odor coding in the LH to be tuned along a variety of parameters. The specific odor-tuning of PNs supports their participation with the coding of distinct “odor scenes” [52,96], and thus 5-HT may differentially affect the processing of specific odor combinations. Furthermore, the LH itself has a zonal organization [21,47,49,100] and the projections of each lv-PN and v-PN into these zones may reflect distinct roles in coding of distinct stimulus features such as valence, identity or intensity [21,30,31,96]. Within the AL, the R24H08 v-PNs densely innervate the DC3 and VL2a glomeruli, both of which respond to plant-derived odors that likely influence reproductive success [69,101,102], and project to the ventral LH which is associated with the processing of pheromones [47,49]. The consistent expression of the 5-HT1A and 1B receptors by these v-PNs implies that serotonin will suppress their activity while simultaneously having a different modulatory effect on other v-PNs that process odors with a distinct behavior context. Although we currently lack an “ecological 5-HT receptor expression logic”, it will be exciting to eventually generate a systematic comparison of the odor space encoded by different v-PNs and lv-PNs and their 5-HT receptor expression profile.

Within cell-class diversity of receptor expression adds a critical layer of nuance to our understanding of the organization of modulation within the AL. It is difficult to determine how many neuron classes are present within a network [103] and heterogeneity within a cell class can blur the lines between functional categories of neurons [104,105]. In our case, the diversity of 5-HT receptor expression patterns within the lv-PNs and v-PNs may merely reflect functional subdivisions within the two populations. The v-PNs consist of at least three morphological classes (uniglomerular, multiglomerular and panglomerular) [39] and there may be other factors such as biophysical properties that could provide functional subdivisions within the v-PNs. Within cell-class diversity is rife within the AL, especially with regards to LNs which vary widely in their morphology, transmitter content, physiological properties, connectivity and odor tuning breadth [23,106–114], so it is not difficult to imagine that similar diversity exists within the v-PNs and lv-PNs. The intrinsic diversity of these groups of AL neurons and the diversity of 5-HT receptors, as well as receptors for other neuromodulators that likely target lv-PNs and v-PNs [58,112], would therefore provide a large parameter space within which odor coding can be fine-tuned within a variety of contexts.

## Acknowledgements

This work was funded by an NIH DC-016293 Award to A.M.D., and a Grant-In-Aid of Research (G20141015669888) from Sigma Xi, The Scientific Research Society to T.R.S.. We also thank Eric Horstick and Kevin Daly for thoughtful comments on the manuscript, Kaleb Hatch and Joshua Taylor for technical assistance, and Dr. Tzumin Lee, the Janelia Research Campus Fly Core and Dr. Hermann Dierick for providing fly stocks.

## Lead contact

Further information and requests for resources and reagents should be directed to and will be fulfilled by the lead contact, Andrew M. Dacks (Andrew.Dacks mail.wvu.edu).

## Materials availability

*Drosophila* stocks generated in this paper are available from the lead contact without restriction upon request.

## Author Contributions

Conceptualization: J.J., M.M. and A.M.D.; Methodology: J.J., M.M., K.L.R. and A.M.D.; Formal analysis: J.J. and M.M.; Investigation: J.J., M.M., T.R.S., J.D.R., F.S., E.J.F., D.E.M. and A.M.D.; Writing–Original Draft: J.J., M.M., T.R.S. and A.M.D.; Writing–Review & Editing: J.J., M.M., T.R.S.; Visualization: J.J., M.M., T.R.S., J.D.R., E.J.F., D.E.M. and A.M.D.; Supervision: A.M.D.; Funding Acquisition: A.M.D.

## Declaration of interests

The authors have no competing interests.

---

**Table 1.**
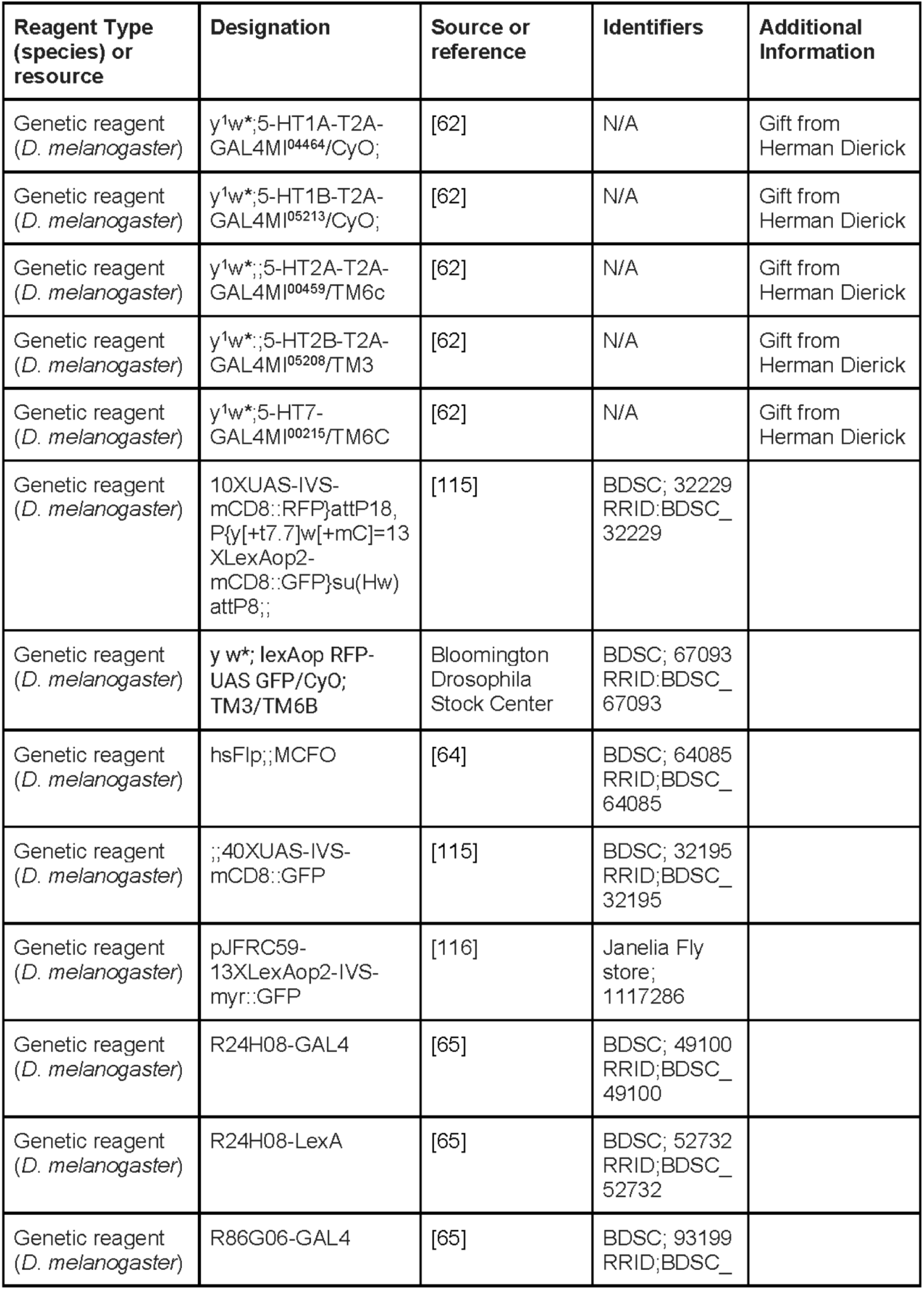
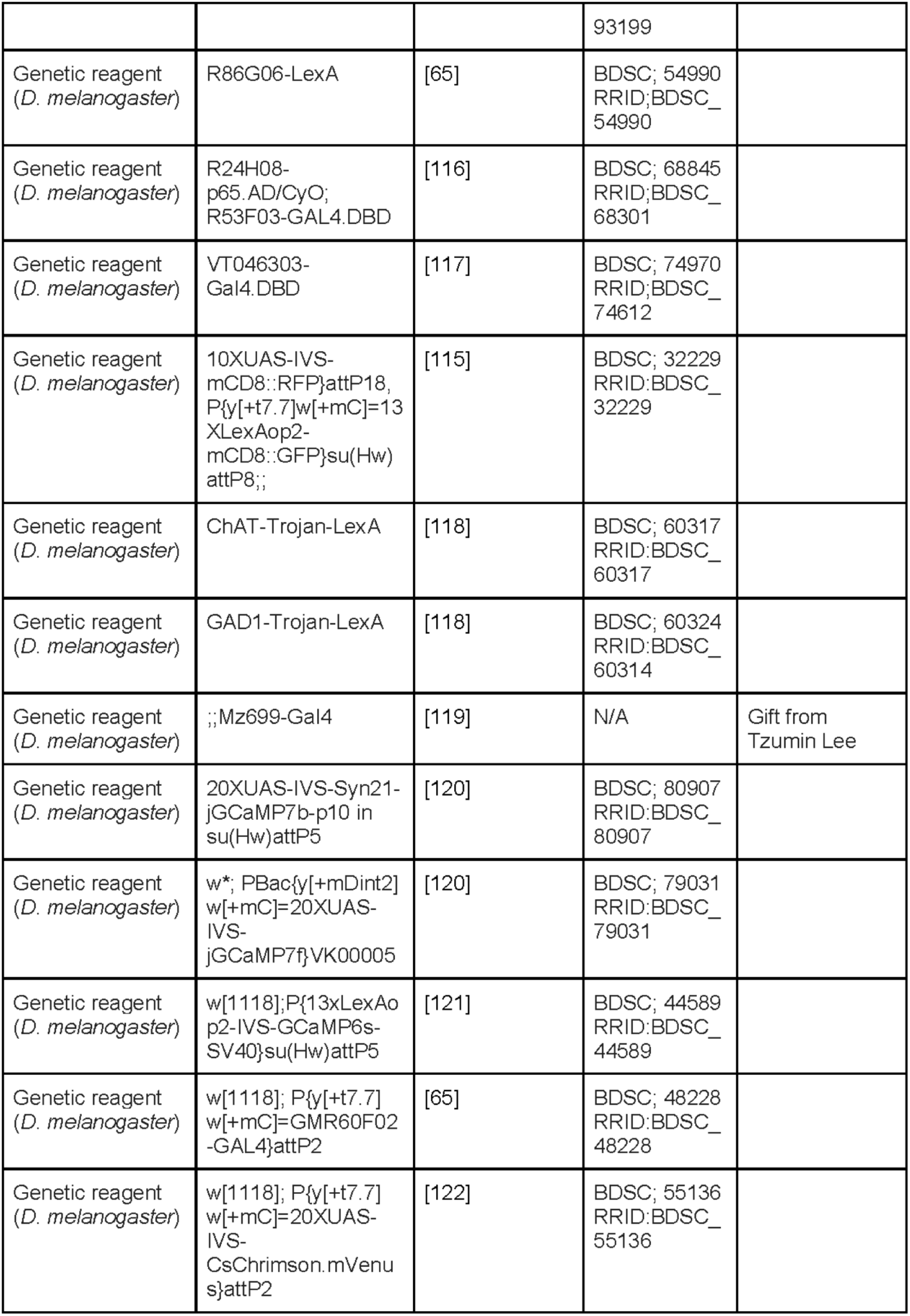
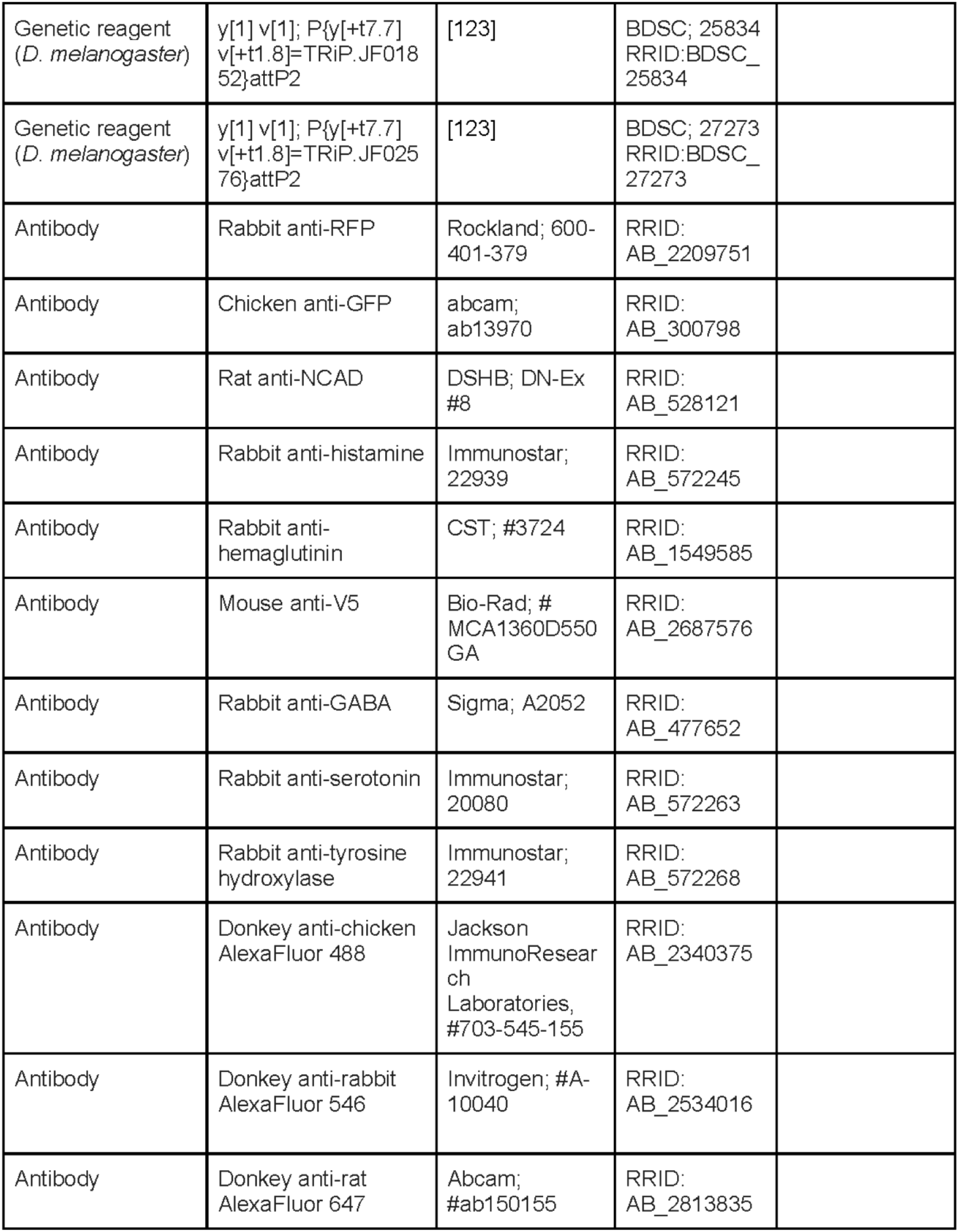
Key Resources and Reagents

**Table 2:**
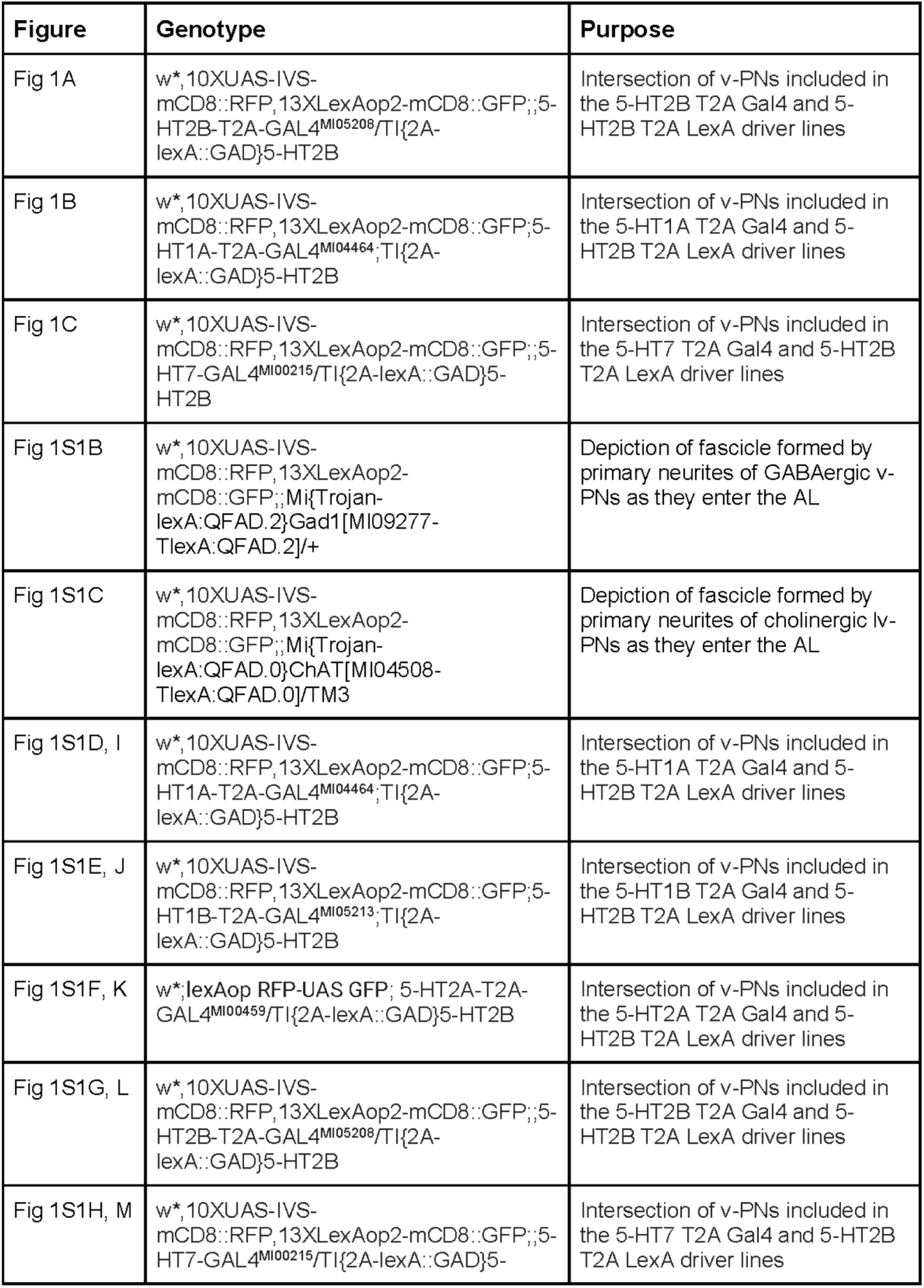
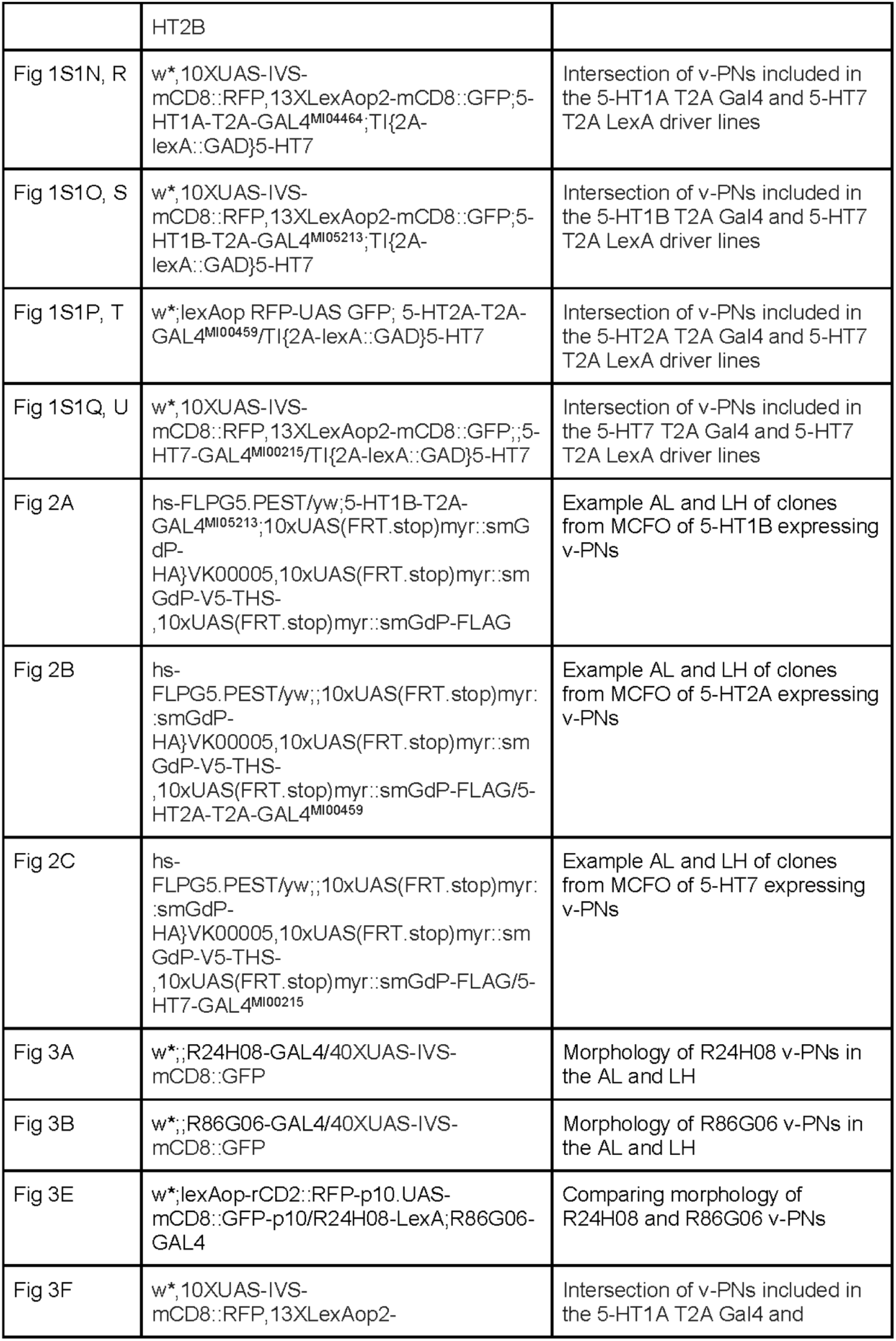
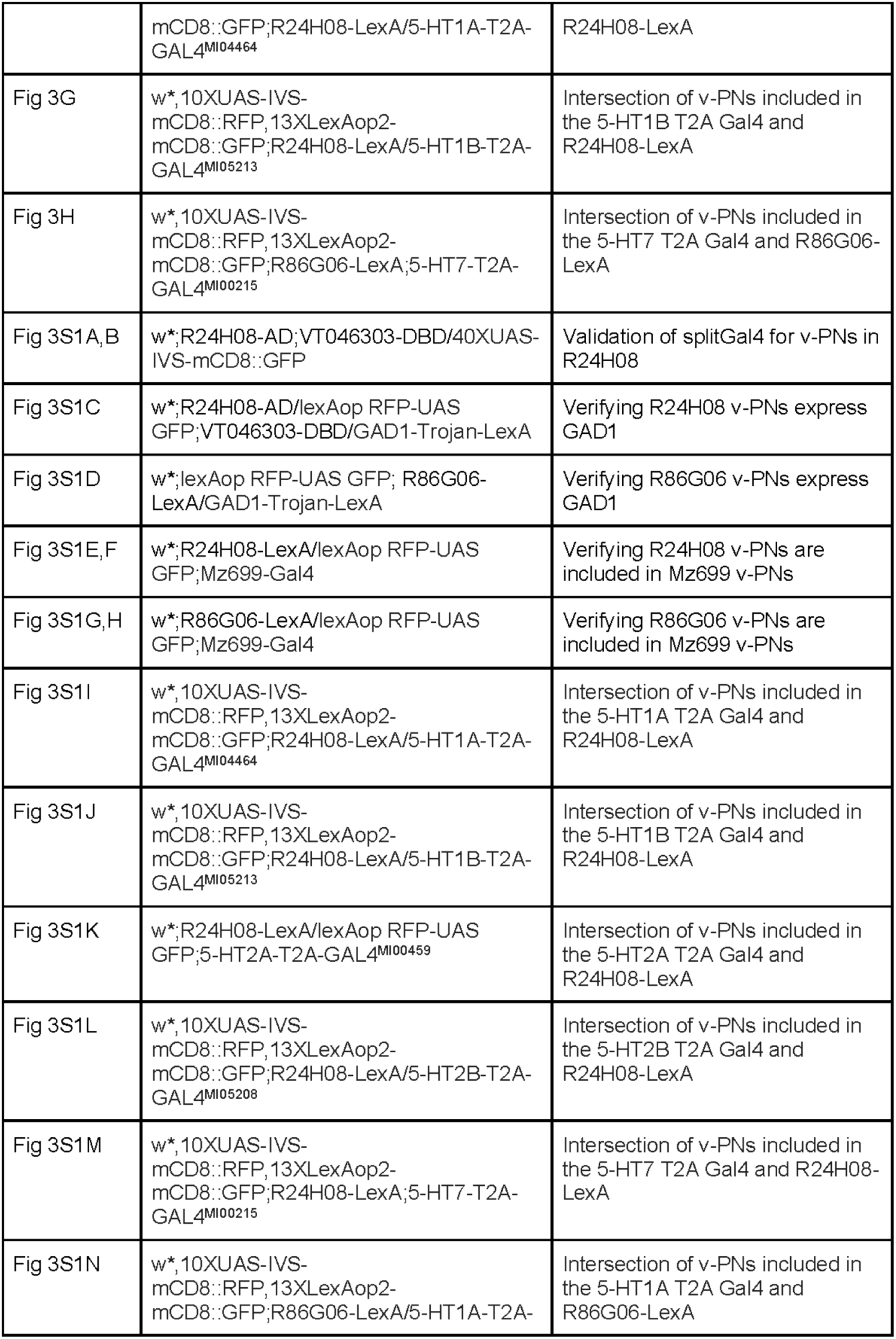
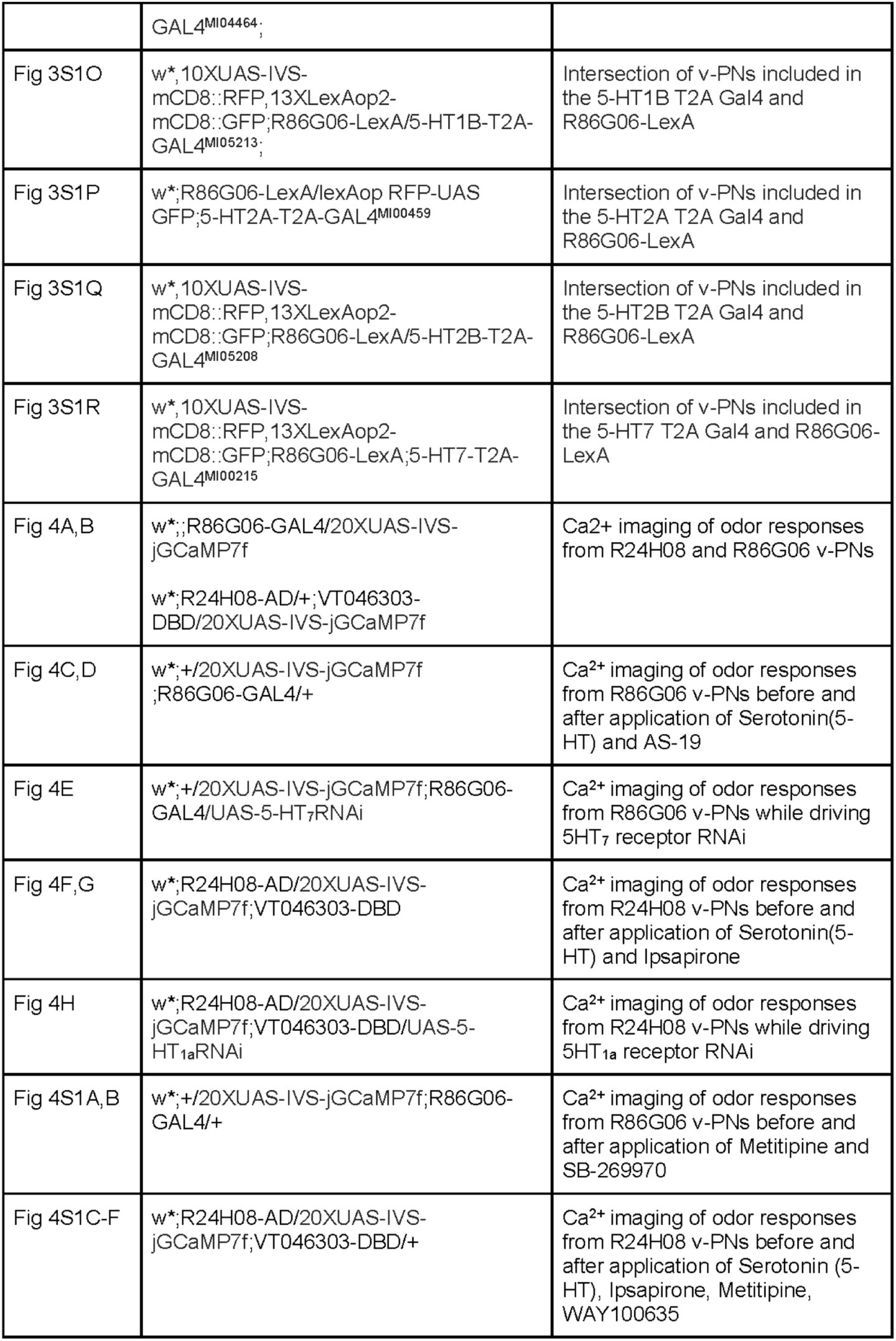
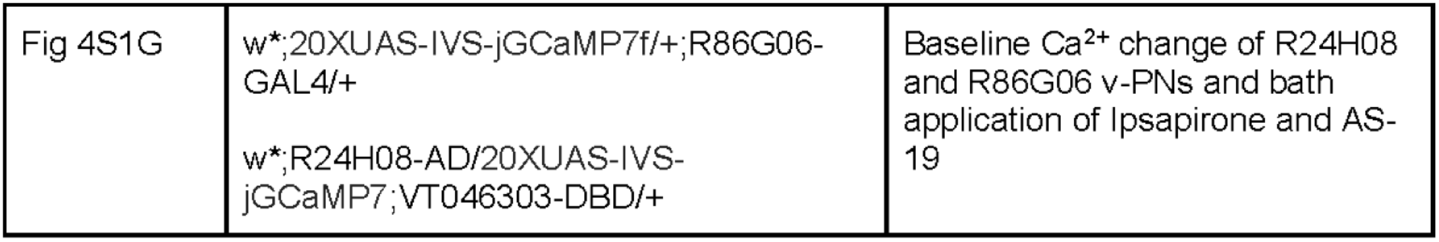
Genotype of flies in each figure

## Methods

### Fly stocks

All fly stocks were raised on a standard cornmeal/agar/yeast medium at 24°C on a 12:12 light/dark cycle at ∼60% humidity.

### Immunocytochemistry and image acquisition

Immunocytochemistry and image acquisition was performed as described previously [19,59]. Briefly, brains were dissected in *Drosophila* External Saline (CSHL recipe) for 30 minutes and placed in 4% paraformaldehyde solution at 40°C, washed in PBS with 0.5% Triton-X (PBST), incubated in blocking solution which consisted of 4% IgG free BSA (Jackson ImmunoResearch, CAS:001-000-162) in PBST and incubated for 48 hours in primary antibodies in 4% BSA in PBST with 5mM sodium azide (PBSAT). Brains were then washed and blocked as above, and incubated for 48 hours in secondary antibodies in 4% BSA in PBSAT. Finally, brains were washed twice with PBST, twice with PBS, run through an ascending glycerol series (40%, 60%, and 80% glycerol in water respectively) and mounted in VectaShield (Vector Labs Burlingame, CA #H-1000). Brains were scanned using an Olympus confocal microscope FV1000 equipped with 40x silicon oil immersion lens. Images were viewed and analyzed using Olympus FluoView software and processed using Inkscape and CorelDRAW vector quality graphics software.

For MCFO experiments, animals were exposed to different intervals of heat-shock to obtain sparse expression of v-PN clones from each 5-HT T2A-Gal4 line (0 min for 5-HT1A, 5-HT1B and 5-HT2A, 0, 5 and 10 min for 5-HT2B and 10 min for 5-HT7) and were dissected 2-7 days later. Due to strong 5-HT2B-T2A-Gal4 expression in ORNs [19] and 5-HT7-T2A-Gal4 in Johnston’s Organ neurons (data not shown), antennal and maxillary palp ablations were conducted two days before heat-shock to prevent obstruction of v-PN processes in the AL or v-PN cell bodies near the AMMC, respectively. MCFO scans could include either only lv-PNs, only v-PNs, or both and if multiple v-PNs and/or vl-PNs were flipped out, their glomerular innervation patterns were grouped for consideration in one sample. We used the coupon collector’s problem [124,125] to determine the number of ALs to image based on v-PN counts previously reported for each 5-HT receptor [19]. Any samples in which AL neurons other than v-PNs and lv-PNs were labeled were discarded to avoid misattributing glomerular innervation. The v-PNs and lv-PNs could be distinguished as lv-PN cell bodies enter the AL through an “arched-shape” fascicle that is located posterior to the “S-shaped” fascicle through which the v-PNs enter the AL [23,95]. These fascicles were also used to identify v-PNs and lv-PNs for the co-expression experiments in **Figure 1** and the fascicles are visible using reporters lines for their respective transmitters; a glutamic acid decarboxylase 1 (GAD1) Trojan T2A-LexA and a choline acetyltransferase (ChAT) Trojan T2A-Gal4 line [119] **(Figure S1A-C)**.

Scans of brains for the MCFO analysis were selected for inclusion if they contain unobstructed views of v-PN cell bodies with their respective morphology and clear glomeruli innervation that could be traced back to an AL fascicle and exiting tract (mALT for lv-PN and mlALT for v-PN). We can only claim that this approach included v-PNs that were most likely to be expressed using MCFO and likely does not represent the total population of v-PNs that express a given 5-HT receptor. We looked at innervation patterns across 47 glomeruli (DA4l and DA4m, VM5d and VM5v and VAlm and VA1l were each merged), comparing across several established glomerular maps [23,126–128]. If more than one v-PN or lv-PN could be visualized in a preparation, the glomerular innervation for all neurons were included for the single sample, rather than trying to distinguish the innervation patterns of the individual neurons.

### Hierarchical Clustering

To resolve whether v-PNs that express distinct 5-HT receptors innervate distinct glomeruli, we hierarchically clustered individual 5-HT receptor expressing v-PNs by the complement of AL glomeruli each v-PN innervates as in [129]. Briefly, individual 5-HT receptor expressing v-PNs were hierarchically clustered using Ward’s method (“ward.D2”) and Euclidean distance using the “dist”, “hclust”, and “as.dendrogram” functions of the base-R *stats* package. The optimal number of k-clusters for this dendrogram were computed using two distinct methods: (1) calculating the maximal average silhouette width using the “find_k” function of the *dendextend* package; and, (2) calculating the gap statistic using the “NbClust” function of the *NbClust* package. Both methods identified that two k-clusters were optimal for the underlying data being hierarchically clustered.

### Pharmacology

Pharmacological agents and their concentrations used in this study: 5-HT receptor agonist (10^-5^M serotonin hydrochloride, TCI Chemicals, CAS:153-98-0) was made fresh every day before the start of experiments and the aliquot was shielded from light, 5-HT receptor antagonist (10^-5^M methiothepine mesylate salt, Sigma, CAS:74611-28-2), 5-HT1A receptor antagonist (10^-5^M WAY-100635 maleate, Abcam, CAS:634908-75-1), 5-HT1A receptor agonist (10^-5^M Ipsapirone, Rndsystems, CAS:95847-70-4), 5-HT7 receptor antagonist (10^-5^M Sb 258719, Tocris, CAS:1217674-10-6), 5-HT7 receptor agonist (10^-5^M AS-19, Tocris, CAS:1000578-26-6). The stock solutions of these drugs were diluted in extracellular physiological saline which contained: 103 mM NaCl, 3 mM KCl, 5 mM TES, 8 mM Trehalose, 10 mM Glucose, 26 mM NaHCO_3_, 1 mM NaH_2_PO_4_, 1.5 mM CaCl_2_, 4 mM MgCl_2_, pH was then adjusted to 7.2 with NaOH.

### Fly preparation for *in vivo* calcium imaging and odor delivery

All *in vivo* calcium imaging experiments were performed using a custom built (Scientifica, Clarksburg, USA) 2-photon microscope system and Mai Tai HP Ti Sapphire laser (Spectra-Physics, Milpitas, CA). Data was acquired using Retiga R6 Microscope Camera (QImaging, Surrey, Canada) and ScanImage acquisition software (v.5.5, Vidrio Technologies). All recordings were taken at a frame rate of 3.4Hz. Both male and female flies were used in experiments. Flies were first anesthetized on ice and then placed on the recording dish containing a square aluminum foil sheet (10mm x 12mm) in the center of a plastic dish with an imaging window (∼1mm x 1mm) sized to affix a fly. Once the fly was securely positioned it was then permanently fixed using LED-UV plastic welder kit (BONDIC, SK8024, NY). Once the fly was glued in place with the head fixed so that the antennae remained dry saline application, then a small incision was made using 26-gauge needles (BD PrecisionGlide Needle, 305110-26g, NJ) and covering tissue was then removed in order to expose the dorsal side of the brain. Odorants used in experiments: 1-hexanol (Sigma Aldrich, cat. no.471402), 1-octen-3-ol (Sigma Aldrich, cat. no. O5284), benzaldehyde (Sigma Aldrich, cat. no. B1334), ACV (Heinz), Farnesol (Sigma Aldrich, cat. no. F203), Orange Peel, acetic acid (Sigma Aldrich, cat. no. A6283), acetophenine (Sigma Aldrich, cat. no. A10701), 1:100 dilution was used for all odors and diluted in mineral oil (Sigma Aldrich, cat. no. M5904). Odors were delivered similarly to [130]. Briefly, odorant dilutions were pipetted onto pieces of Whatman filter paper in 5cc glass syringes with 20-gauge needles inserted through a rubber septum (Thermogreen LB-2 Septa, 20633, Bellefonte, PA) into a common air stream directed at the antennae. Common and odor air streams originated from compressed air that was first carbon filtered, then re-humidified before being split to the constant airflow line (2.5L/min regulated using a Dwyer VFA-25-BV flowmeter) and the airflow (.8L/min regulator using a Dwyer VFA-23-BV flowmeter) delivered to a solenoid (Parker, 001-0028-900, Hollis, NH) that could switch between an empty cartridge and an odor cartridge. Custom MatLab script (Matlab version 2018b, provided by Kaleb Hatch) was used to send a 5V TTL pulse to a 50W power source (CUI Inc, 102-3295-ND, Tualatin, OR) to actuate the solenoid. Constant airflow was directed to the antennae via a central glass tube with two ports holding rubber septa into which the empty cartridge and odor cartridge could be inserted to introduce a second airstream at a 45° angle. Odorants were delivered by activating the solenoid to switch the second airflow from the empty cartridge to the odor cartridge for 2-3 times depending on experimental protocol for 1 second. Raw imaging data was then imported into FIJI and then ROIs were drawn. Extracted ROI fluorescence signal data was then processed using a custom Matlab Script which normalized to baseline fluorescence signals (F, fluorescence averaged across 3 seconds before the first odor stimulation) and data was visualized as percent change in fluorescence from average values (ΔF/F), script then produced representation of percent change in fluorescence as a Heatmap (Matlab version 2018b, script provided by Keshav L. Ramachandra) or average ΔF/F odor responses were then processed in GraphPad for statistical analysis.

### Statistical analyses

Number of animals used in each experiment were between 8 and 12, (number of animals and trials are stated in figure legends). Statistical analysis was carried out using GraphPad Prism software (GraphPad Prism version 8.0, 2018). Paired sample T-tests were performed in all GCaMP imaging datasets and passed Shapiro-Wilk normality tests.

### Connectomic analyses

Connectomic analyses in **Figure 3** were performed using the Hemibrain v.1.2.1 dataset [66] accessed via neuPrintExplorer (https://neuprint.janelia.org/). Within this dataset, nine v-PNs were identified as candidates for the v-PNs expressed by the R24H08 and R86G06 LexA driver lines. These were identified based on their glomerular innervation and their projection patterns within the lateral horn, and the number of v-PNs with these characteristics were consistent with the number of cells observed within each driver line (five for R24H08-LexA and four for R86G06-LexA). The body IDs for the five v-PNs similar to those in R24H08 were 5813077810, 1857143769, 948834414, 1889883818, and 1415825344 and the body ID for those v-PNs similar to the four v-PNs in R86G06 were 791298858, 760264077, 729219639, and 698526273. To determine the postsynaptic partners of each v-PN, the downstream partners were queried for each individual v-PN and an arbitrary threshold of three synapses was set for inclusion. Postsynaptic targets that were an “orphan” or not ascribed to a cell class within neuPrint were also excluded from analysis. To determine the degree to which v-PNs from R24H08 and R86G06 converged upon the same postsynaptic partners, the “common connectivity” query was used for all nine v-PNs simultaneously. An arbitrary threshold of three synapses was set for inclusion and the resultant connectivity table was exported to generate graph plots using CytoScape v3.9.1.

---

**Figure S1.**
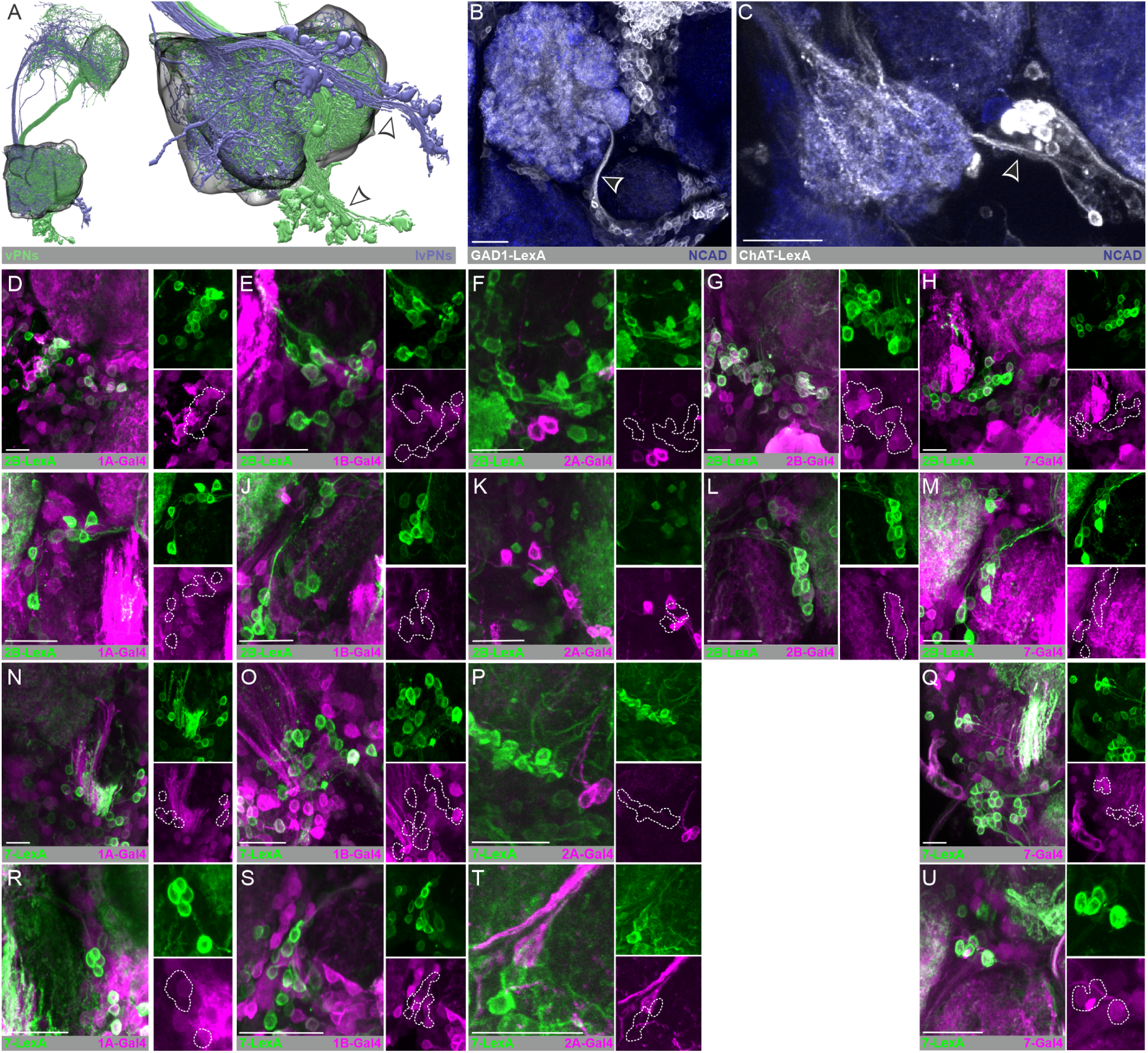
Additional representative images of 5-HT receptor co-expression. **(A)** Volumetric reconstructions of v-PNs (green) and lv-PNs (blue) from the Hemibrain EM dataset highlighting the different tracts (arrowheads) formed by the primary processes of each cell cluster. **(B)** The fascicle formed by the cholinergic v-PNs is highlighted by a GAD1-T2A-LexA driver (white). Neuropil delineated by N-cadherin (blue). **(C)** The fascicle formed by the cholinergic lv-PNs is highlighted by a ChAT-T2A-LexA driver (white). Neuropil delineated by N-cadherin (blue). **(D-U)** Example images of intersection between 5-HT receptor T2A-Gal4 (magenta) and LexA (green) drivers. **(D-H)** Example images from v-PNs of intersection between the 5-HT2B-T2A-LexA and **(D)** 5-HT1A-T2A-Gal4 (15 ALs from 9 brains), **(E)** 5-HT1B-T2A-Gal4 (14 ALs from 8 brains), **(F)** 5-HT2A-T2A-Gal4 (17 ALs from 12 brains), **(G)** 5-HT2B-T2A-Gal4 (14 ALs from 7 brains) and **(H)** 5-HT7-T2A-Gal4 (18 ALs from 9 brains). **(I-M)** Example images from lv-PNs of intersection between the 5-HT2B-T2A-LexA (Average cell body count = 10.97±1.04, n = 73 AL) and **(I)** 5-HT1A-T2A-Gal4 (12 ALs from 6 brains), **(J)** 5-HT1B-T2A-Gal4 (16 ALs from 8 brains), **(K)** 5-HT2A-T2A-Gal4 (17 ALs from 12 brains), **(L)** 5-HT2B-T2A-Gal4 (13 ALs from 7 brains) and **(M)** 5-HT7-T2A-Gal4 (17 ALs from 9 brains). **(N-Q)** Example images from v-PNs of intersection between the 5-HT7-T2A-LexA and **(N)** 5-HT1A-T2A-Gal4 (19 ALs from 10 brains), **(O)** 5-HT1B-T2A-Gal4 (16 ALs from 8 brains), **(P)** 5-HT2A-T2A-Gal4 (16 ALs from 22 brains) and **(Q)** 5-HT7-T2A-Gal4 (13 ALs from 7 brains). **(R-U)** Example images from lv-PNs of intersection between the 5-HT7-T2A-LexA (Average cell body count = 5.97±0.48, n = 50 ALs) and **(R)** 5-HT1A-T2A-Gal4 (12 ALs from 9 brains), **(S)** 5-HT1B-T2A-Gal4 (12 ALs from 8 brains), **(T)** 5-HT2A-T2A-Gal4 (15 ALs from 11 brains) and **(U)** 5-HT7-T2A-Gal4 (11 ALs from 7 brains). All scale bars are 20um.

**Figure S2.**
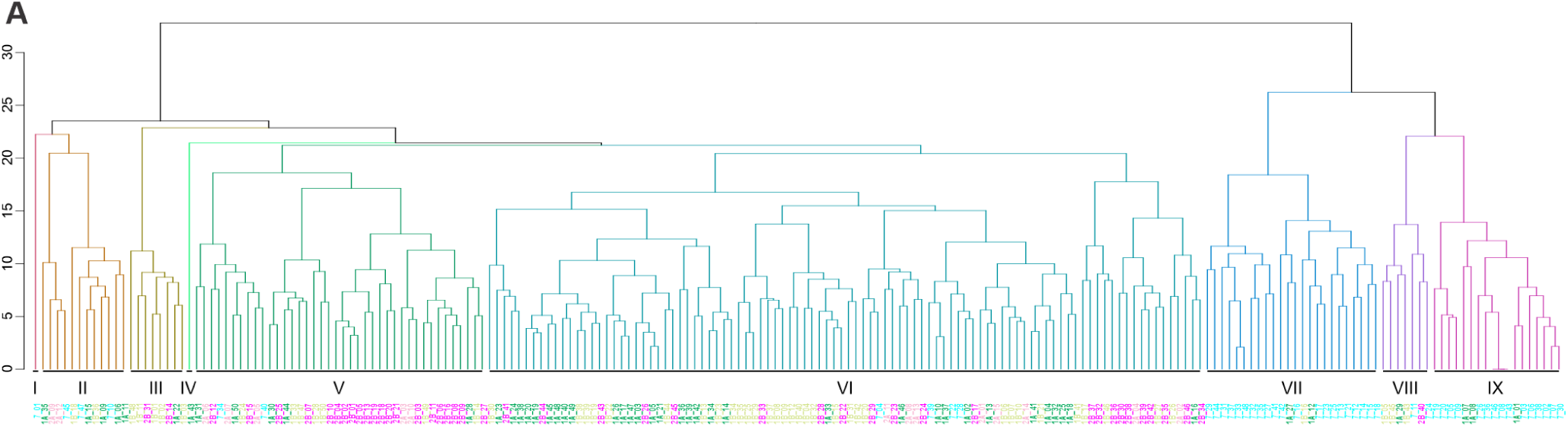
Hierarchical clustering of 5-HT receptor T2A MCFO glomerular innervation patterns. **(A)** Hierarchical clustering analysis of the glomerular innervation patterns of glomeruli innervated by v-PNs and lv-PNs stochastically labeled via MCFO for each 5-HT receptor T2A-Gal4 line. A total of nine clusters (I-IX, each with their own color) were obtained and clones within each cluster are colored based on the MiMIC T2A-Gal4 from which they derive (green; 5-HT1A, yellow; 5-HT1B, salmon; 5-HT2A, pink; 5-HT2B, cyan; 5-HT7).

**Figure S3.**
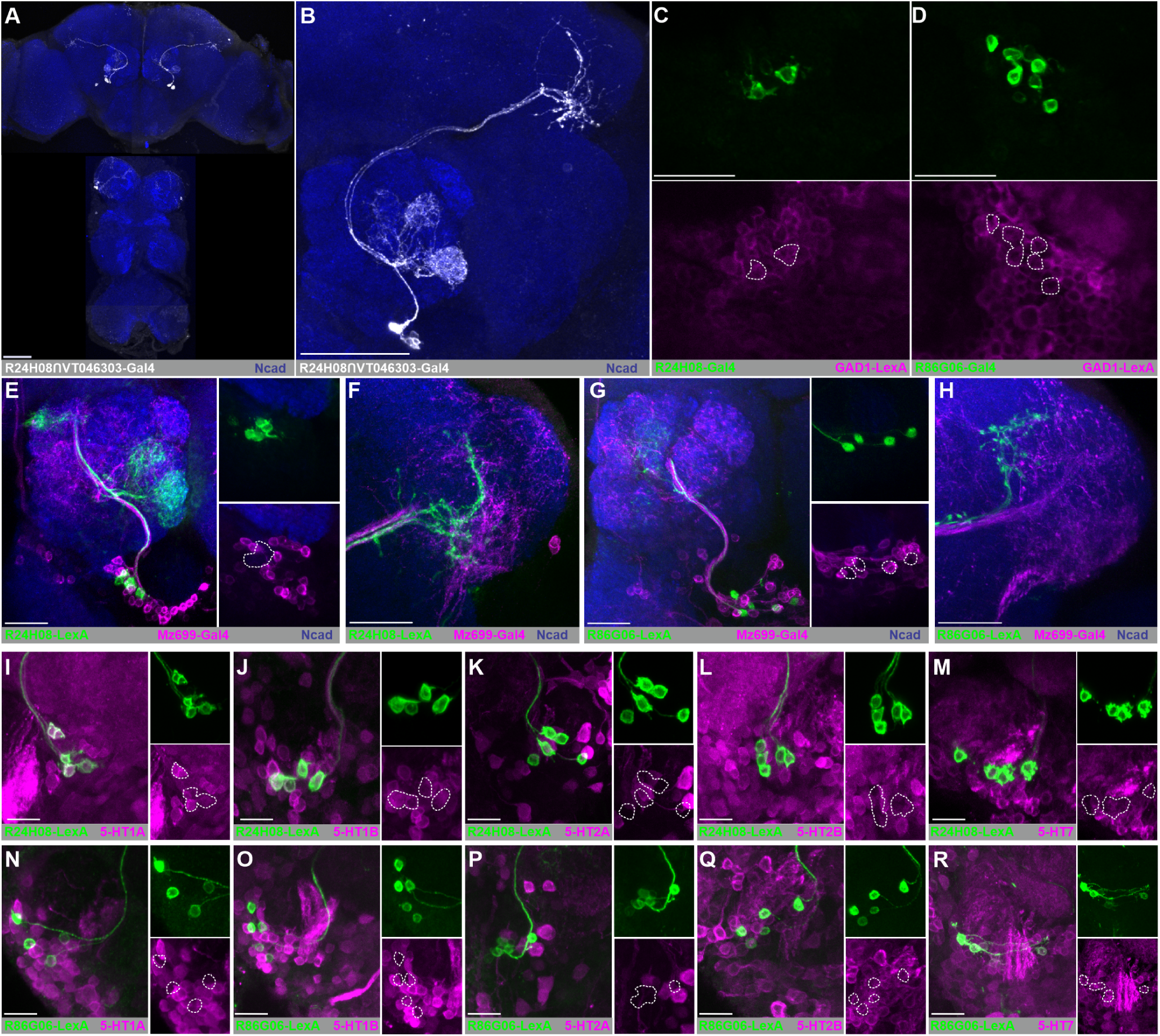
Additional images of v-PNs expressed in the R24H08 and R86G06 driver lines. **(A)** Expression pattern of the R24H08 *∩* VT046303 split-Gal4 (white) in the brain and ventral nerve cord. NCAD serves as a neuropil marker (blue). **(B)** Expression pattern of the R24H08 *∩* VT046303 split-Gal4 AL and LH. **(C-D)** The v-PNs in both **(C)** R24H08-Gal4 and **(D)** R86G06-Gal4 overlap with expression of a GAD1-LexA. **(E-H)** The v-PNs in **(E-F)** R24H08-LexA and **(G-H)** R86G06-LexA (green) are not included in the v-PNs present in the Mz699-Gal4 (magenta) driver line. **(E & G)** Projections of v-PNs in **(E)** R24H08-LexA and **(G)** R86G06-LexA combined with the v-PNs from Mz699-Gal4 within the AL. **(F & H)** Projections of v-PNs in **(F)** R24H08-LexA and **(H)** R86G06-LexA combined with the v-PNs from Mz699-Gal4 within the LH. **(I-M)** Intersection between the R24H08-LexA and T2A-Gal4 lines for the **(I)** 5-HT1A, **(J)** 5-HT1B, **(K)** 5-HT2A, **(L)** 5-HT2B and **(M)** 5-HT7 receptors. **(N-R)** Intersection between the R86G06-LexA (green) and T2A-Gal4 lines (magenta) for the **(N)** 5-HT1A, **(O)** 5-HT1B, **(P)** 5-HT2A, **(Q)** 5-HT2B and **(R)** 5-HT7 receptors. All scale bars = 20um.

**Figure S4.**
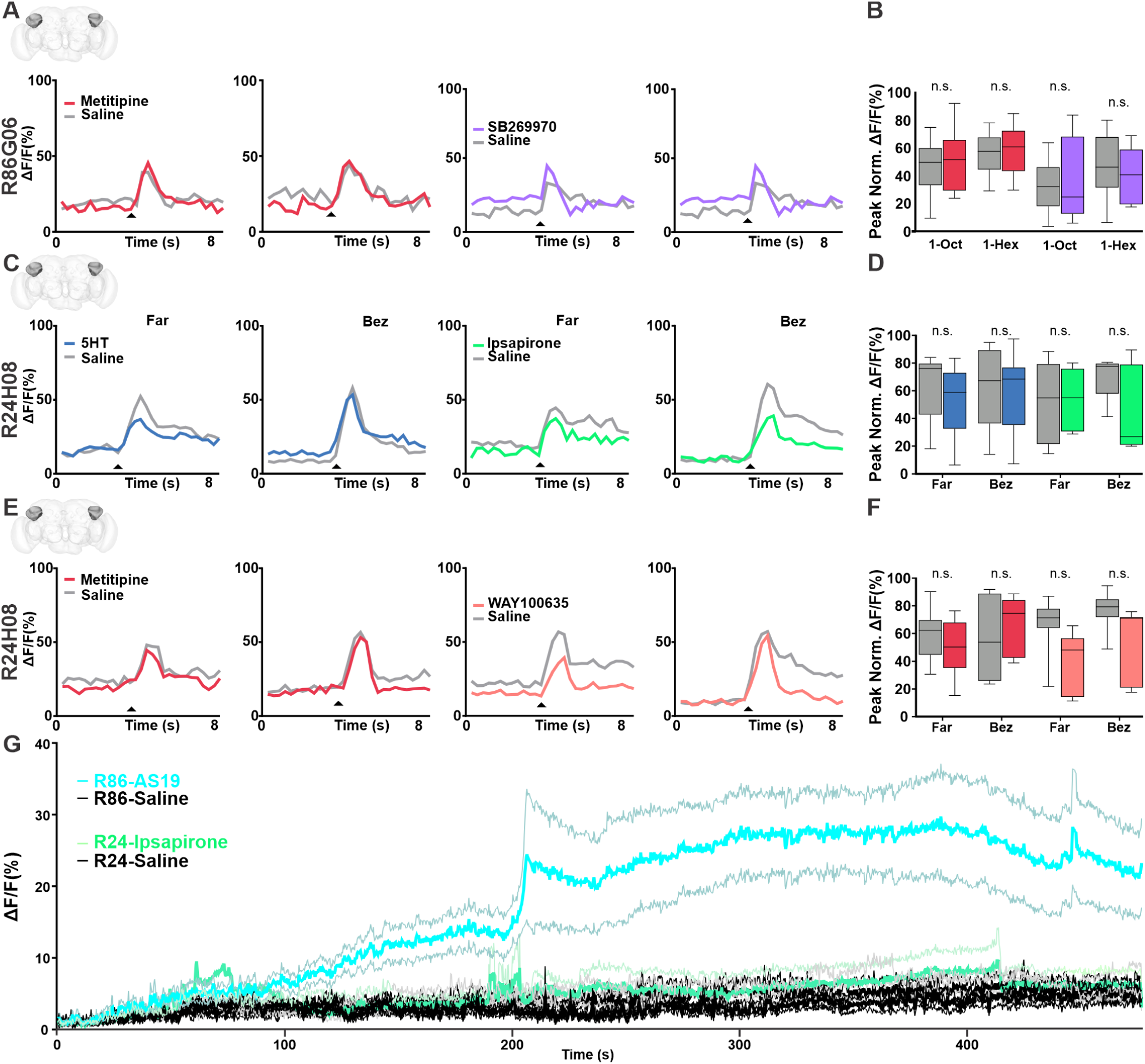
Additional 5-HT receptor pharmacology experiments. **(A)** Average responses of R86G06 v-PNs to 1-octen-3-ol (1-Oct) or 1-hexanol (1-Hex) before (gray traces) and after application of the pan-5-HT receptor antagonist metitipine (Red) and the the 5-HT7 receptor antagonist SB269970 (purple). **(B)** Neither metitipine nor SB269970 affected odor-evoked responses of R86G06 v-PNs. Metitipine, n=11 (1-Oct), n=11 (1-Hex); SB269970, n=10 (1-Oct), n=11 (1-Hex) animals, 3 repeats, ns; paired sample t-test. **(C)** Average responses of R24H08 v-PNs recorded from processes in the lateral horn to farnesol (Far) or benzaldehyde (Bez) before (gray traces) and after application of 5-HT (blue) and the 5-HT1A/1B receptor agonist ipsapirone (green) and the the 5-HT7 receptor antagonist WAY100635 (orange). **(D)** Neither 5-HT nor ipsapirone affected odor-evoked responses of R24H08 v-PNs recorded within the lateral horn. 5-HT, n= 11 (Far), n=12 (Bez); Ipsapirone, n=8 (Far), n=7 (Bez) animals, 3 repeats, ns; paired sample t-test. **(E)** Average responses of R24H08 v-PNs recorded from processes in the lateral horn to farnesol (Far) or benzaldehyde (Bez) before (gray traces) and after application of the pan-5-HT receptor antagonist metitipine (Red) and the the 5-HT1A receptor antagonist WAY100635 (orange). **(F)** Neither metitipine nor WAY100635 affected odor evoked responses of R24H08 v-PNs recorded within the lateral horn. Metitipine, n=6 (Far), n=6 (Bez); WAY100635, n=7 (Far), n=7 (Bez) animals, 3 repeats, ns; paired sample t-test. **(G)** Average response of R86G06 v-PNs and R24H08 v-PNs to bath application of AS-19 and ipsapirone respectively over 8 minutes. N= 7-12 samples.

## Notes

### Competing Interest Statement

The authors have declared no competing interest.

